# HSP90 facilitates oncogenic alterations of metabolism in B-cell lymphomas

**DOI:** 10.1101/2020.05.29.119925

**Authors:** M. Nieves Calvo-Vidal, Nahuel Zamponi, Jan Krumsiek, Max A. Stockslager, Maria V. Revuelta, Jude M. Phillip, Rossella Marullo, Nikita Kotlov, Jayeshkumar Patel, Shao Ning Yang, Lucy Yang, Tony Taldone, Catherine Thieblemont, John P. Leonard, Peter Martin, Giorgio Inghirami, Gabriela Chiosis, Scott R. Manalis, Leandro Cerchietti

## Abstract

HSP90 is critical for maintenance of the cellular proteostasis. In cancer cells, HSP90 also becomes a nucleating site for the stabilization of multiprotein complexes including signaling pathways and transcription complexes. Here, we described a novel role of HSP90 in the cytosolic compartmentalization of metabolic pathways in proliferating cancer cells. We found that HSP90 assists in the organization of metabolic enzymes into non-membrane-bound functional compartments termed metabosomes. Under experimental conditions that conserved the cellular proteostasis, we demonstrated that the compartmentalizing activity of HSP90 is critical to sustain the coordinated synthesis of multiple metabolites required for energy production, maintenance of the cellular biomass and secretion of immunometabolites. Conversely, inhibition of the nucleating capacity of HSP90 modified the topology of cytosolic metabosomes before protein degradation was apparent decreasing the efficiency of MYC-driven metabolic pathways. Inhibition of HSP90 decreases cancer metabolism in B-cell lymphoma cells and patients providing a novel mechanism of activity for this class of drugs.

## INTRODUCTION

Diffuse large B-cell lymphoma (DLBCL), the most common non-Hodgkin B-cell lymphoma, and Burkitt lymphoma (BL) are highly proliferative diseases that exhibit, accordingly, an increased rate of catabolic glucose and glutamine metabolism[1]. DLBCL and BL cells augment their aerobic glycolysis rates to produce biosynthetic intermediates for biomass accumulation and secondarily to supplement bioenergetic needs, since the majority of ATP is produced elsewhere, in the mitochondria[2]. This and other metabolic traits of aggressive B-cell lymphomas are determined by the activation of molecular mediators including MYC and MTOR/PI3K, among others[3–5]. The mechanisms that facilitate the metabolic reprogramming effect of these oncogenic drivers, and thus confer a suvival advantage, have not been fully explained.

Although the expression of alternative isoforms and the increased expression of glycolytic enzymes have been proposed as ways to achieve the high rates of glycolysis observed in cancer cells[6], the role of metabolic enzymes compartments is less understood[7]. Compartmenting metabolic complexes allows the regulation of macromolecular crowding, which modulates protein folding, aggregation and diffusion[8, 9]. Regulated protein crowding is beneficial to activate metabolic pathways that are not constitutively present and/or require protein up-regulation to be established[10]. This has been recognized in proliferating cells by the binding of the glycolytic enzymes glyceraldehyde-3-phosphate dehydrogenase (GAPDH), phosphofructokinase (PFK1), enolase (ENO), pyruvate kinase (PK), lactate dehydrogenase (LDH) and aldolase (ALDO) as well as enzymes of the purine biosynthesis to the cytoskeleton[11–14]. Physical and chemical coordination of molecules is inherently linked to metabolic efficiency as demonstrated by the evolutionary-conserved tendency of multi-enzymatic complex formation[8, 15, 16]. It has been estimated that up to 80% of the metabolic intermediates of a cell serve one purpose[17] such that diffusion into the cytosol will be energetically disadvantageous. Hence, in addition to greater regulatory control, multi-enzymatic complexes that limit substrate diffusion represent a kinetic advantage to cells by increasing the solvation capacity within the cytosol and optimizing funneling of metabolites[8, 15].

Proliferating cells endure the bioenergetic challenge of maintaining cell homeostasis while facilitating biomass accumulation that enables cell growth and division. Metabolic demands are increased even further in cancer cells as a consequence of the stress imposed by an aberrant biological background and unstable microenvironmental conditions, affecting both proliferating and non-proliferating cells[1, 6]. To increase the efficiency of the cellular metabolism, stressors promote the regulated crowding of metabolic proteins. For instance, under nutrient or hypoxic stress certain glycolytic enzymes coalesce into cytosolic clusters or “glycosomes” providing higher rates of glycolysis[18, 19]. In human cells cultured under purine metabolic stress, multiple enzymes catalyzing *de novo* purine biosynthesis colocalize to intracellular foci known as “purinosomes” to maintain the purines output[20–22]. Moreover, inosine monophosphate dehydrogenase 2 (IMPDH2) of the purine biosynthesis pathway physically associates with cytidine triphosphate synthetase (CTPS) of the pyrimidine biosynthesis pathway in mammalian cells grown under specific amino acid starvation[23]. Although some enzymatic polymers can be self-regulated, others require the assistance of stress chaperones such as the formation of hypoxic stress glycosomes in yeast[18] and the assembly of purinosomes in transformed human cells[24, 25].

The chaperome, an assembly of molecular stress chaperones and their many partners, assist in protein folding and in reducing protein aggregation to maintain cellular proteostasis. In cancer cells the heat shock protein 90 (HSP90) chaperome can be incorporated with the heat shock protein 70 (HSP70) chaperome to assemble higher-order structures regarded as “epichaperomes”[26]. Epichaperomes can increase, not just balancing, the fitness of the proteome in cancer cells by assisting in the formation of multi-protein complexes[26, 27]. This function becomes critical for cancer proliferation on a background of internal and external stress that results from biological instability and changing microenvironmental conditions[6]. Cancer cells are not only metabolically stressed by oncogene expression but also from the activation of detoxification processes and the secretion of macromolecules and metabolites to the microenvironment[1].

In this study, we elucidated the role of the HSP90 epichaperome in the organization of metabolic enzymes into cytosolic non-membrane-bound functional compartments in B-cell lymphoma cells. We identified cytosolic metabolic proteins interacting with active HSP90 epichaperome in native proteostasis conditions without requiring exogenous protein expression. We then integrate this data in a global scale with the metabolic alterations occurring in lymphoma cells after short-term administration of an HSP90 inhibitor. We described specific associations of metabolic enzymes and metabolites in functional units termed metabosomes. We finally demonstrated that this novel metabolic function of HSP90 is required to fulfill biomass, energetic and metabolic secretory requirements of B-cell lymphomas and, specifically, to support the metabolic program driven by MYC. Conversely, HSP90 inhibition disaggregated metabosomes ultimately decreasing B-cell lymphoma metabolism in pre-clinical models and patients.

## RESULTS

### Oncogenic HSP90 nucleates multi-protein complexes containing metabolic enzymes

To study the HSP90 protein interactome in lymphoma cells, we took advantage of a method that we previously demonstrated as able to maintain the biochemical and thermodynamic stability of active HSP90-containing epichaperome complexes[27–29]. We characterized the endogenous HSP90 interactome in cytoplasmic lysates of the DLBCL cell lines OCI-LY1 and OCI-LY7 by using a chemical bait that captures HSP90-containing epichaperome and its associated interacting proteins, followed by identification by mass spectrometry[30] (**Fig. 1A**). To ensure maximal capture of HSP90 complexes, we performed these experiments under conditions of excess bait and by adding molybdate to induce HSP90 oligomerization and stabilize the dynamic HSP90 multi-protein complexes under normal cellular proteostasis. This methodology does not require exogenous protein expression while isolating multiprotein complexes in their native functional configuration. Pathway enrichment analysis of the cytoplasmic HSP90 interactome revealed a significant overrepresentation of proteins from cellular processes typically involving multiprotein megacomplexes such as signaling, protein degradation, RNA translation and metabolism (**Fig. 1A, Table S1**). We and others have previously described the role of HSP90 in maintaining the active conformation of the BCR signalosome, transcriptional complexes and polyribosomes[29–31]; however, the role of cytoplasmic HSP90 in cellular metabolism has not been addressed.

**Figure 1.**
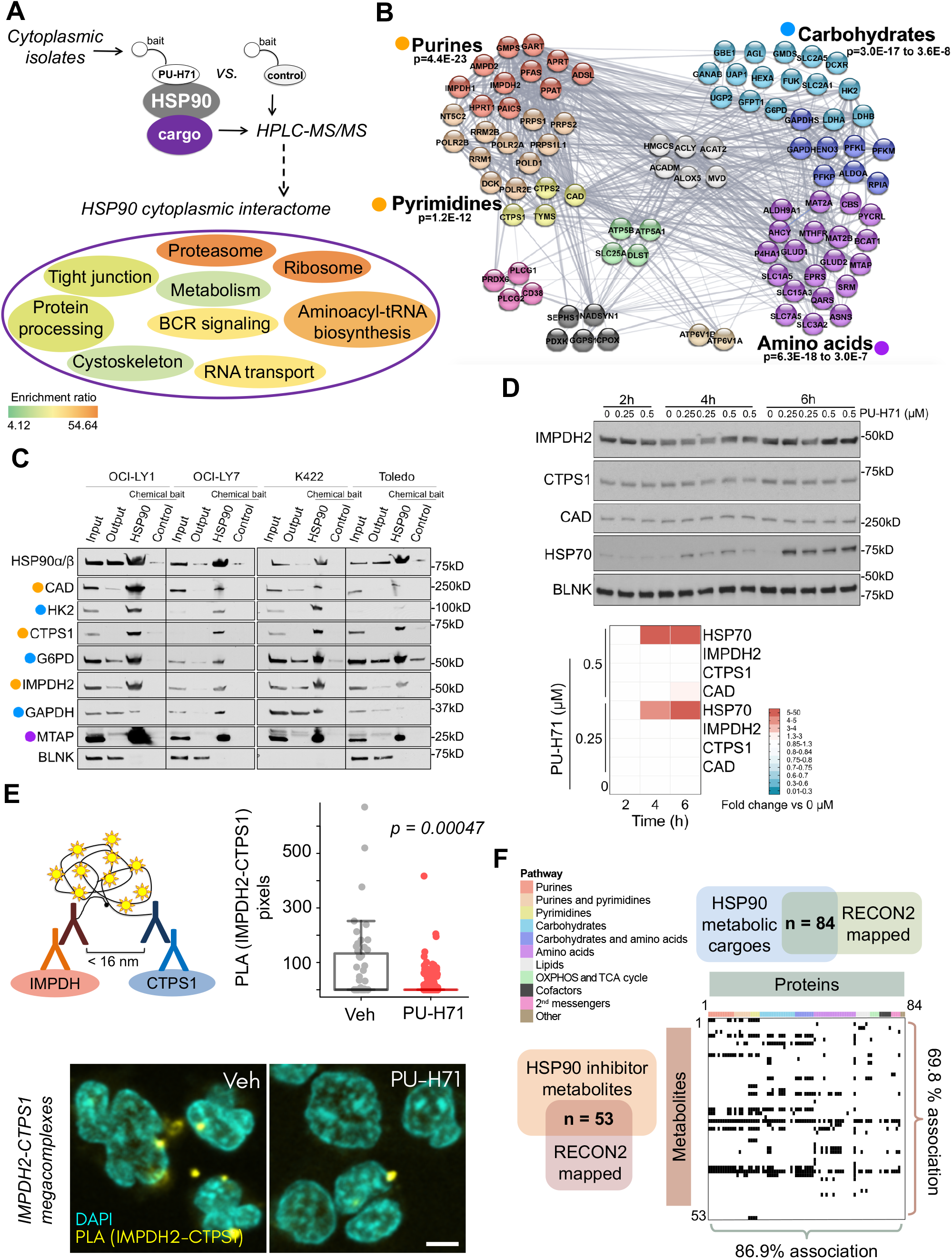
Functional association of the HSP90 metabolic interactome and the metabolome into metabosomes. **A:** Pathways enriched in the active HSP90 interactome from the cytoplasmic fraction of OCI-LY1 and OCI-LY7 DLBCL cells. Enrichment is indicated by the color bar, and the relative significance (adjusted p value) by the font size. **B:** STRING representation of the metabolic pathways from the HSP90 interactome. The significance of the enrichment is under each metabolic category. **C:** Validation of representative enzymes from the HSP90 interactome belonging to the three enriched pathways (from B, as indicated by colored dots) in the DLBCL cell lines OCI-LY1, OCI-LY7, Karpas 422 and Toledo. The lanes represent original lysates (input), proteins remaining after HSP90 chemical affinity purification (output), HSP90 chemical affinity purification (HSP90) and inert chemical affinity purification (control). HSP90 was used as positive control and BLNK was used as negative control. **D:** Time course of the abundance of HSP90 metabolic interactome components IMPDH2, CTPS1 and CAD upon HSP90 inhibition with increasing concentrations of PU-H71 (0.25 and 0.5 μM) or vehicle (0) in OCI-LY7 cells. HSP70 was used as molecular readout of effective HSP90 inhibition and BLNK as a non-HSP90-cargo control. Densitometry of blots are shown at the bottom as color-coded fold changes over vehicle control. **E:** Representative imaging of IMPDH2 and CTPS1 endogenous megacomplexes (yellow puncta) in OCI-LY1 DLBCL cells treated with vehicle or PU-H71 0.5 μM for 6 h. The bar represents 5 μm. Quantification is shown on top. **F:** Association of the subset of HSP90 metabolic cargoes mapped to Recon 2 (n = 84) with the subset of differentially changed metabolites upon PU-H71 treatment mapped to Recon 2 (n = 53). Proteins are ordered and color-coded by pathways. Matched protein-metabolite pairs are shown as black rectangles in the association plot.

To further categorize the subset of metabolic proteins (hereon the HSP90 metabolic interactome) we used STRING (Search Tool for the Retrieval of Interacting Genes). The resulting functional association network showed clusters significantly enriched for enzymes involved in the metabolism of nucleotides (purines and pyrimidines), carbohydrates and amino acids (**Fig. 1B, Table S2**). To independently validate proteins as part of the HSP90 metabolic interactome we used the DLBCL cell lines OCI-LY1, OCI-LY7, Karpas422 and Toledo, and, by chemical affinity purification followed by immunoblotting, we confirmed several enzymes from these pathways including G6PD (glucose 6-phosphate dehydrogenase), GAPDH (glyceraldehyde-3-phosphate dehydrogenase) and HK2 (hexokinase 2), from “carbohydrates”; CAD (carbomyl-phosphate synthetase, aspartate transcarbamylase and dihydroorotase), IMPDH2 (inosine monophosphate dehydrogenase 2) and CTPS 1 (cytidine 5-triphosphate synthetase 1), from “nucleotides”; and MTAP (methylthioadenosine phosphorylase), from “amino acids” (**Fig. 1C**). The cytosolic protein BLNK (B-cell linker) was used as a negative control (**Fig. 1C**).

To determine whether the HSP90 metabolic interactome was constituted of particularly thermally unstable proteins, we compared their general structural and biochemical properties[32] obtained from protein databases. There were no common features in this subset of proteins (**Fig. S1**), and neither an enrichment in proteins classified as unstable as determined by their melting temperature[32] (**Fig. S1**). This suggests that this is not a subset of particularly unstable proteins. In accordance with this notion, HSP90 inhibition using PU-H71 at anti-neoplastic doses of 200 and 500 nM for up to 6 h did not result in decreased abundance of the HSP90 metabolic interactome components CAD, IMPDH2, and CTPS1 (**Fig. 1D**). In contrast, between 14 to 24 h of HSP90 inhibition was necessary to observe the typical decrease of these cargo proteins (**Fig. S1**). However, up-regulation of HSP70 demonstrated a complete inhibition of HSP90 as early as 4 h after PU-H71 administration (**Fig. 1D**). In these experiments, BLNK served as a non HSP90-cargo control (**Fig. 1D**).

Further analysis of the reactions catalyzed by components of the HSP90 metabolic interactome indicated that these are more likely to be key steps, defined as irreversible, committed and/or rate-limiting reactions. Specifically, the HSP90 metabolic interactome is responsible for seven out of eight of the regulatory steps in the purines and pyrimidines biosynthesis pathway: PPAT (phosphoribosyl pyrophosphate aminotransferase), IMPDH1/2, CAD, CTPS1/2, TS (thymidylate synthetase), RRM1/2 (ribonucleotide reductase subunits 1 and 2) and RRM2B (ribonucleotide reductase regulatory subunit M2B) (**Fig. S2**). Additionally, the highly regulated PRPS1/2 is also an HSP90 cargo. Similarly, five of the six key enzymes in the “carbohydrates” pathway are part of the HSP90 metabolic interactome: HK2, G6PD, PFK1, GFPT1 (glutamine-fructose 6-phosphate transaminase) and UGP2 (UDP-glucose pyrophosphorylase) (**Fig. S2**). In yeast and mammalian cells these enzymes have been shown to be part of multi-enzymatic complexes such as the glycosome and purinosome[8]. Because these are nonmembrane-bound proteins, our data suggests that HSP90 may be facilitating the spatial organization of these enzymes in the cytosol of DLBCL cells.

### HSP90 topologically organizes its metabolic interactome into metabosomes

Under high nucleotide demand conditions, the enzymes IMPDH2 of the purine biosynthesis and CTPS1 of the pyrimidine biosynthesis form homo- and heteropolymers to increase their efficiency and maintain the synthesis of purines and pyrimidines nucleotides[23]. The molecular mechanism that facilitates these complexes has not been fully elucidated. Because these two enzymes are components of the HSP90 metabolic interactome (**Fig. 1B-C**), we investigated the role of HSP90 in the cytosolic topology of IMPDH2-CTPS1 multi-enzymatic metabolic megacomplexes. We treated OCI-LY1 cells with the HSP90 inhibitor PU-H71 (500 nM) for 6 h and assessed the endogenous interaction of IMPDH2 and CTPS1 by a protein proximity ligation assay. We found that HSP90 inhibition altered the cytosolic topology of IMPDH2-CTPS1 complexes demonstrated by a significant decrease (p = 0.00047) in the number of stable interactions (**Fig. 1E**).

To determine the functional consequences of inhibiting the HSP90 metabolic interactome at the cellular scale, we integrated the HSP90 metabolic interactome with cellular metabolomics performed in the same DLBCL cell lines, OCI-LY1 and OCI-LY7, 6 h after the administration of 500 nM PU-H71 (vs. vehicle control). We found 90 metabolites significantly changed upon PU-H71 administration (**Fig. S3**). To establish the degree of functional association between these metabolites and proteins we developed a metabolic reconstruction model where metabolites are linked to enzymatic reactions by adapting the data from RECON2[33]. We considered any positive direct association between a metabolite and enzyme when they share the reaction and/or metabolic pathway. We found that 70% of the metabolites that significantly changed with PU-H71 were associated with at least one protein of the HSP90 metabolic interactome (**Fig. 1F**); conversely, 87% of the components of the HSP90 metabolic interactome associated with at least one PU-H71-changed metabolite (**Fig. 1F**), indicating a high degree of functional association. The full extent of the integrated proteomics and metabolomics data was visualized by overlaying it onto the KEGG human metabolism map (**Fig. S3**). The high correlation between the metabolic interactome and their associated metabolites indicates that HSP90 functionally organizes the cytosol of lymphoma cells into multiple “metabosomes”.

### HSP90 inhibition decreases nutrient utilization in DLBCL cells

To determine the metabolic advantage provided by active HSP90 in the cytosol of lymphoma cells, we evaluated the effects of shortterm HSP90 inhibition on nutrient utilization. Central to maintenance of the biomass in cancer cells is the oxidation of glucose via glycolysis. We measured the glucose uptake from the culture medium and the secretion of lactate as the end-product of glycolysis into the culture medium, in OCI-LY1 and OCI-LY7 cells treated with 200 nM and 500 nM PU-H71 or vehicle for 6 and 14 h. We found a decrease in glucose uptake (**Fig. 2A**) and lactate secretion (**Fig. 2B**) in both cell lines. As expected, due to an accumulative effect, the differences were more significant after 14 h. This data indicate HSP90 inhibition impairs glycolysis. We independently confirmed this effect by measuring the extracellular acidification rate (ECAR) in the same cell lines (**Figs. 2C** and **S4**). The basal ECAR was significantly lower in both cell lines (**Fig. 2D**). Upon addition of oligomycin to inhibit the mitochondrial respiration and elicit the maximal ECAR, cells treated with PU-H71 could not reach the maximal ECAR of vehicle-treated cells (**Fig. 2D**), leading to a significant increase in the glycolytic reserve by 15-20% (**Fig. 2D**). This suggests that lymphoma cells with active HSP90-dependent metabosomes have increased efficacy to metabolize glucose.

**Figure 2.**
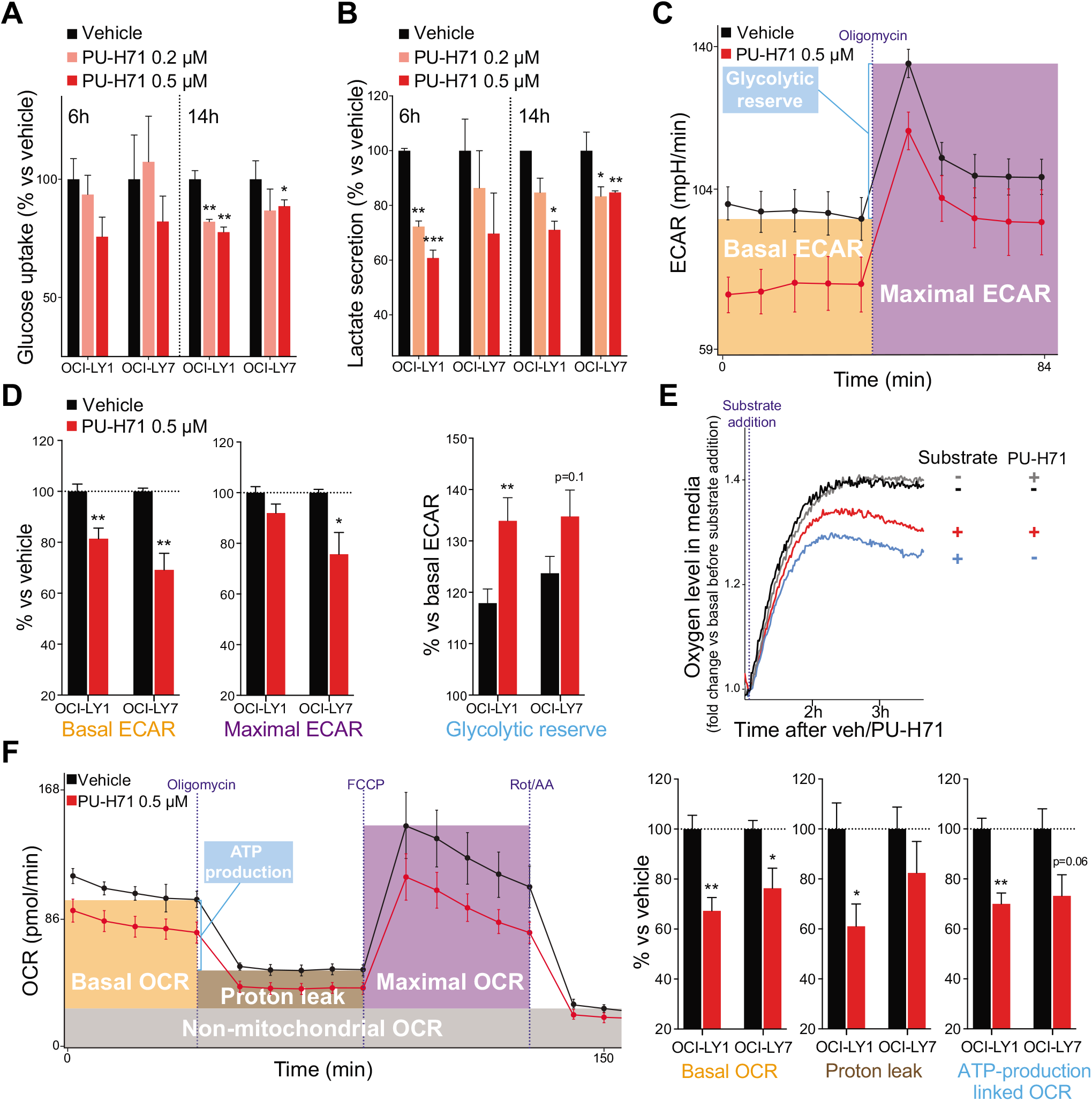
HSP90 inhibition decreases nutrient utilization. **A-B:** Glucose uptake (A) and lactate excretion (B) in OCI-LY1 and OCI-LY7 cells treated with vehicle (black), PU-H71 0.2 μM (pink) and PU-H71 0.5 μM (red) for 6 h and 14 h. Data is normalized to vehicle-treated cells. **C:** Extracellular acidification rate (ECAR) of OCI-LY1 cells treated with vehicle (black) or PU-H71 (red), at baseline and upon oligomycin treatment to estimate the maximal ECAR. Error bars are SD of 10 replicate wells. Representative experiment of triplicates shown. **D:** Mean basal ECAR, maximal ECAR and glycolytic reserve capacity in OCI-LY1 and OCI-LY7 cells treated with vehicle (black) or PU-H71 (red), normalized to vehicle. **E:** Mitochondrial respiration as determined by real-time measurement of oxygen levels in the tissue culture medium of OCI-LY1 cells in absence of glutamine with vehicle (black) or PU-H71 (grey), and presence of glutamine with vehicle (blue) or PU-H71 (red). Representative experiment of triplicates shown. **F:** Oxygen consumption rate (OCR) of OCI-LY1 cells treated with vehicle (black) or PU-H71 (red) at baseline, upon oligomycin treatment to determine proton leak and OCR-linked ATP production, upon FCCP to estimate maximal OCR, and upon rotenone/antimycin A to estimate non-mitochondrial OCR. On the right, mean basal OCR, proton leak and ATP production linked to OCR in OCI-LY1 and OCI-LY7 cells treated with vehicle (black) or PU-H71 (red). In all panels, unless stated differently, error bars are SEM of 3 independent experiments. p values were calculated by T-test. n.s., not significant, *<0.05, **<0.01 and ***<0.001.

In addition to glucose, DLBCL cells consume glutamine in the mitochondria via glutaminolysis, a process that feeds the tricarboxylic acid cycle and readily increases mitochondrial respiration. To determine whether the inhibition of cytosolic HSP90 could have an impact on glutaminolysis and respiration, we measured oxygen consumption of OCI-LY1 cells exposed to PU-H71. For this, we monitored in realtime the oxygen levels in the culture medium immediately after addition of glutamine as substrate. At this time point, glutamine uptake was not affected by PU-H71 (**Fig. S4**). As expected, the addition of glutamine induces oxygen consumption, indicating the cells are capable of glutaminolysis (**Fig. 2E**); however, in presence of PU-H71 this effect is less pronounced (**Fig. 2E**). Similar results were obtained with the Burkitt lymphoma cell line Raji (**Fig. S4**). We independently confirmed the effects of PU-H71 on mitochondrial respiration by measuring the oxygen consumption rate (OCR) in DLBCL cells cultured with glucose, glutamine and pyruvate. We found that PU-H71 decreased basal OCR in OCI-LY1 and OCI-LY7 cells (**Fig. 2F** and **S4**), as well as respiratory parameters such as the proton leak and the ATP-production linked OCR (**Fig. 2F**). There were no differences in the maximal OCR or non-mitochondrial OCR between treated and non-treated cells (**Fig. S4**). These data indicate that cytoplasmic HSP90 activity supports both glycolysis and, secondarily, mitochondrial respiration. The mechanism for this latter effect likely involves a more efficient channeling of substrates provided by the organization of cytosolic metabosomes since PU-H71 does not penetrate the mitochondria[34].

### HSP90 inhibition decreases macromolecule biosynthesis and biomass gain

Whereas in non-proliferating cells respiration is mainly an ATP-producing catabolic process, in proliferating cells respiration serves also as a crucial anabolic role by providing both direct and indirect intermediates, as well as cofactors, for various processes including synthesis of proteins, lipids and nucleotides. In this regard, the dihydroorotate dehydrogenase (DHODH) step of the pyrimidines biosynthesis is coupled to the mitochondrial electron transport chain and respiration. The de novo synthesis of nucleotides is also contingent on the ribose 5-phosphate formed primarily by the pentose phosphate pathway (PPP). Pathway mapping of the metabolic changes caused by HSP90 inhibition in OCI-LY1 and OCI-LY7 cells shows that key precursors orotate and ribose 5-phosphate are decreased upon HSP90 inhibition (**Fig. S5**). Furthermore, glucose carbon (D-glucose-^13^C^6^) tracing in OCI-LY1 cells indicates that under normal conditions where the oncogenic form of HSP90 is active, the canonical oxidative and non-oxidative phase of the PPP are active and produce ribose 5-phosphate (and NAPDH) for nucleotide synthesis (**Fig. 3A**). Upon HSP90 inhibition (PU-H71, 6h), the production of ribose 5-phosphate decreases while of ribulose 5-phosphate increases (**Fig. 3A**), potentially reflecting a lower activity of the isomerase RPIA, a member of the HSP90 metabolic interactome (**Fig. S2**). Under HSP90 inhibitory conditions a higher proportion of the ribose 5-phosphate is funneled into glycolytic intermediates (**Fig. 3A**). Overall, our data showed that with an active HSP90 a relatively higher proportion of the glucose is oxidized to ribose 5-phosphate likely due to the favored activity of the HSP90 metabolic interactome members G6PD and RPIA (**Fig. S2**).

**Figure 3.**
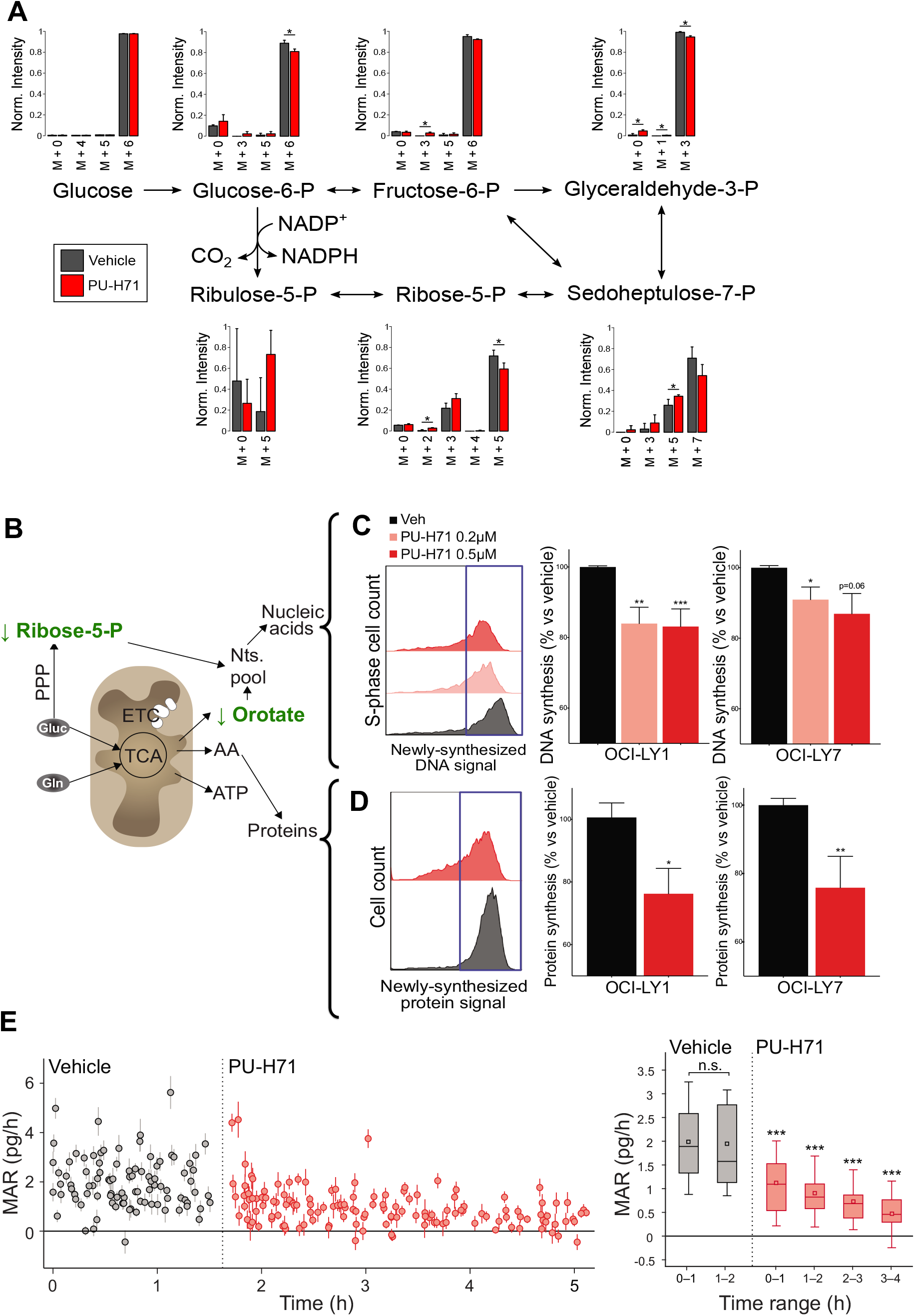
HSP90 inhibition decreases the synthesis of DNA, proteins and biomass gain. **A:** Glucose carbon tracing in OCI-LY1 cells treated with vehicle or the HSP90 inhibitor PU-H71 for 6 h. **B:** Cartoon of the anabolic role of mitochondrial respiration and pentose phosphate pathway. **C:** Detection of newly synthesized DNA by fluorescent thymidine analogue 5-ethynyl-2’-deoxyuridine (EdU) incorporation in OCI-LY1 and OCI-LY7 cells treated with vehicle or PU-H71 0.2 μM and 0.5 μM for 6 h. On the right, mean EdU signal, normalized to vehicle. **D:** Detection of newly synthesized protein by fluorescent amino acid analogue L-homoproparglyglycine (HPG) incorporation in OCI-LY1 and OCI-LY7 cells treated with vehicle or PU-H71 0.5 μM for 6 h. On the right, mean HPG signal, normalized to vehicle. In all panels, error bars are SEM of 3 independent experiments. **E:** Real-time assessment at single-cell resolution of cellular Mass Accumulation Rate (MAR) in OCI-LY1 cells treated with vehicle (black) and upon administration of PU-H71 1 μM to the same culture (red). On the right, mean MAR comparing binned datasets (from time 0 to 1, from 1 to 2, and so on). P values were calculated by T-test. *<0.05, **<0.01 and ***<0.001.

We thus measured the impact of HSP90 inhibition on the synthesis of DNA and proteins by incubating OCI-LY1 and OCI-LY7 cells with thymidine or methionine analogs, respectively, and measuring newly-synthesized DNA and protein, in presence of PU-H71 or vehicle. We found that HSP90 inhibition for 6 h reduced, in both cell lines, DNA synthesis by 10 - 20% and protein synthesis by 25% (**Fig. 3B-D**). Altogether, these results suggest that lymphoma cells with active HSP90-dependent “metabosomes” can more efficiently utilize metabolites required for the synthesis of macromolecules; and effect that may facilitate the maintenance of the cell biosynthetic activity under oncogenic stress. To non-invasively quantify the impact on the cellular biomass with single-cell resolution, we continuously monitored the mass accumulation rate (MAR) of OCI-LY1 cells immediately after exposure to PU-H71 and for up to 4 h, by using a serial array of suspended microchannel resonator[35]. We found a significant reduction of MAR in cells with inhibited HSP90 that occurred as strikingly early as 30 m after PU-H71 administration (**Figs. 3E** and **S5**), denoting a potential interference with nutrient utilization in DLBCL cells upon HSP90 inhibition.

### HSP90 activity supports the production of extracellular immunometabolites

A further implication of our data in DLBCL cell lines is that, in addition to contributing to energy and biomass, certain HSP90-dependent metabolites such as secreted lactate and purines could also have an impact on the tumor micro and macroenvironment. To characterize this effect in the context of the disease, we first determined the most active metabolic pathways in DLBCL by comparing the serum metabolomics (i.e., exo-metabolomics) of 50 DLBCL patients with 25 age- and gender-matched healthy individuals. The exo-metabolomics analysis readily segregated DLBCL patients from healthy individuals as demonstrated by principal component analysis (**Fig. 4A**). We identified 312 significantly different metabolites between these two groups, belonging to several biochemical pathways (**Fig. 4B** and **Table S3**). Among the most overrepresented biochemical categories were nucleotides (e.g., metabolism of purines and pyrimidines), carbohydrates (e.g., pentoses, amino sugars and glycolysis) and aminoacids (e.g., histidine, glutathione, glycine, serine and threonine) (**Fig. 4B**). Remarkably, we found that DLBCL patients present a serum metabolic profile associated with an immunosuppressive microenvironment[36] characterized by increased lactate and pyruvate, decreased tryptophan and increased kynurenine, decreased arginine and increased inosine (**Fig. 4C**).

**Figure 4.**
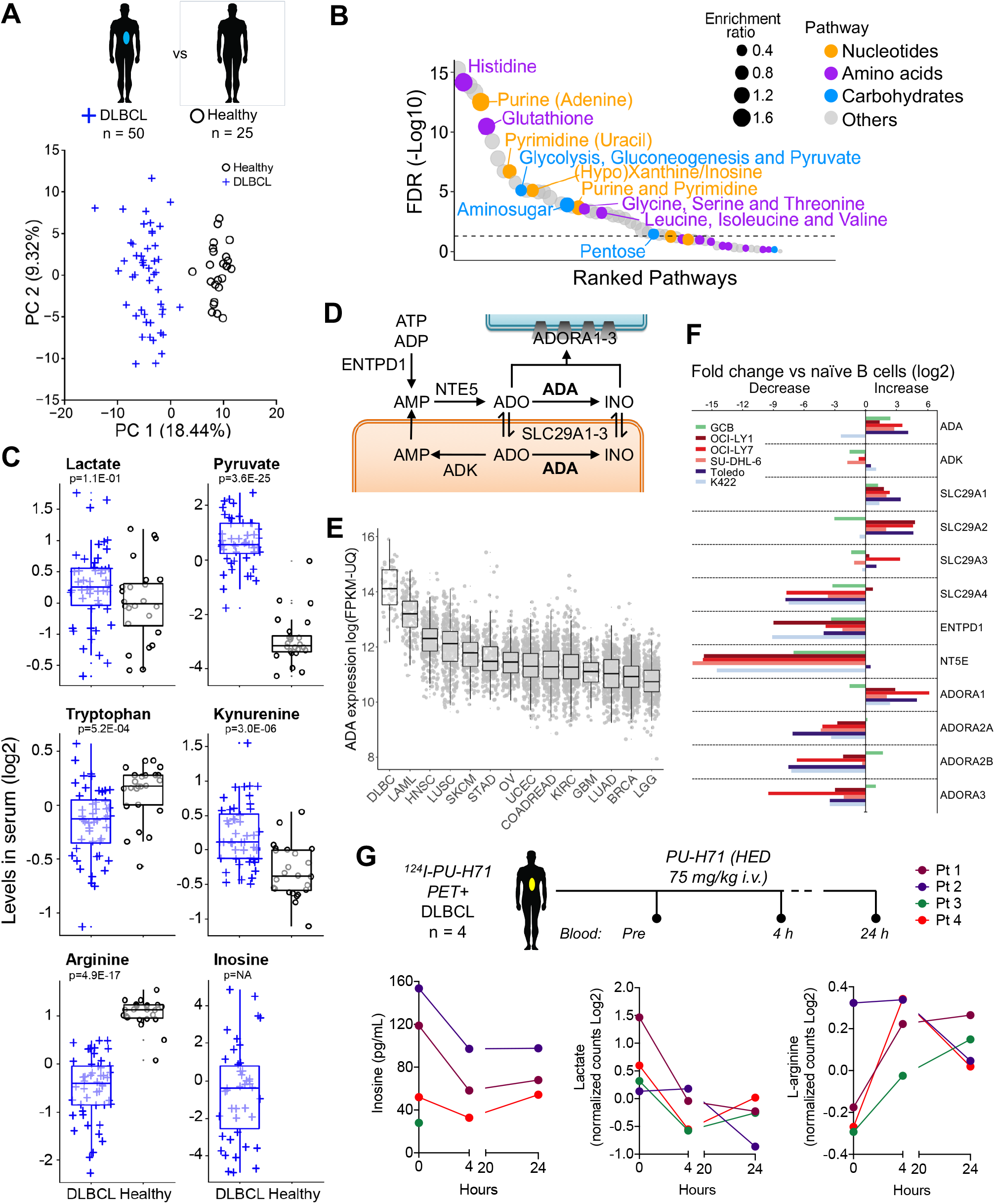
The serum metabolomics of DLBCL patients reflects an immunosuppressive microenvironment characterized by the presence of purines. **A:** Principal component analysis of serum exometabolomics of DLBCL patients (n = 50, blue cross) and age- and gender-matched healthy individuals (n = 25, black circles). **B:** Metabolic pathways significantly different between DLBLC patients and healthy individuals. The relative size of the circles indicates enrichment ratio versus healthy. The color coding indicates super-pathway categorization into “amino acids”, “carbohydrates”, “nucleotides” and “others”. **C:** Box-plots of immune-related metabolites from the serum exo-metabolomics analysis from A. Levels are shown normalized as log2. Adjusted p-values for the comparison are shown below each metabolite. Inosine was not detected in healthy individuals. **D:** Inosine pathway indicating metabolites and intra and extracellular enzymes. **E:** Expression of adenosine deaminase (ADA) in DLBCL vs. other tumors included in the TCGA. **F:** Expression of enzymes and solute transporters involved in the metabolism of inosine in OCI-LY1, OCI-LY7, SU-DHL-6, Karpas422 and Toledo DLBCL cell lines and normal germinal center B-cells (GCB) normalized to levels in naïve B-cells. Representative experiment shown. **G:** Levels of inosine, lactate and L-arginine in 4 patients with epichaperome+ DLBCL (as determined by PET-CT), before and 4 and 24 h after the administration of one dose of PU-H71 i.v.

Inosine represented the most extreme example of these metabolites as it was detected exclusively in DLBCL patients (**Fig. 4C**). Extracellular inosine depends on adenosine levels, which depend mostly on the transport from the intracellular space by nucleoside transporters. Adenosine is deaminated to inosine by adenosine deaminase (ADA) that is present in the intra- and extra-cellular spaces (**Fig. 4D**), and both purines exert similar immune regulatory effects[37]. Extracellular inosine is much more stable than adenosine, which goes in agreement with the lack of detection of adenosine in the serum from our cohorts of healthy or DLBCL individuals (**Table S3**). The prevalence of inosine over adenosine in DLBCL patients could be associated to the higher expression of ADA in comparison with other tumor types (**Fig. 4E**). To further characterize the inosine pathway in DLBCL, we determined the expression of the enzymes, transporters and receptors related to inosine metabolism (**Fig. 4D**) in OCI-LY1, OCI-LY7, SU-DHL-6, Karpas422 and Toledo DLBCL cell lines and found a higher expression of ADA compared to normal human naïve B cells in 4 out of the 5 cell lines analyzed (**Fig. 4F**). Moreover, in three of the DLBCL cell lines the expression was even higher than highly proliferative normal CD77^+^ germinal center B cells (**Fig. 4F**), suggesting that DLBCL cells may have increased capacity of producing inosine. More importantly, lymphoma cell lines overexpressed the adenosine-inosine equilibrating membrane transporters SLC29A1 and SLC29A2 (**Fig. 4F**), responsible for the presence of these purines in the extracellular space. The purinergic receptor ADORA1 was also increased in the lymphoma cell lines (**Fig. 4F**).

To determine the effect of disrupting the nucleotide metabosome in the production of metabolites with immune activity *in vivo*, we measured the immunometabolites inosine, lactate and L-arginine in the plasma of four DLBCL patients with tumors with high ^124^I-PU-H71 uptake by positron emission tomography (PET/CT) that non-invasively indicates the presence of epichaperomes in cancer tissues[27]. These patients from the clinical trial NCT01393509 were administered the human equivalent dose of 75 mg/kg in mice of PU-H71 i.v. and plasma was obtained before and 4 and 24 h after a single dose of the drug. We found that, for each metabolite, 3 out of 4 patients had either a decrease in inosine and/or lactate and/or increase in L-arginine as soon as 4 h after PU-H71 (**Fig. 4G**). In most cases the changes plateau at the 24 h time point (**Fig. 4G**). This finding indicates that immunometabolites are susceptible to low-dose HSP90 inhibition in DLBCL patients. Taken together these data indicate that the activity of the HSP90-dependent metabosomes contributes to sustaining cellular and secretory metabolic needs of DLBCL.

### The HSP90 metabolic interactome supports the metabolic program of MYC in lymphomas

To understand the mechanistic relevance of these findings to the reprogramming of lymphoma metabolism, we analyzed putative transcription factors regulating the expression of the HSP90 metabolic interactome. We found that the promoter of genes coding for the HSP90 metabolic interactome (n = 90, **Fig. 1A**) were significantly enriched for consensus binding sites of MYC and MYC/MAX, SP1, E2F family and ARNT of the HIF1 family (**Fig 5A**), suggesting that these transcription factors may be more reliant on HSP90 to reprogram cell metabolism. Since MYC is known to carry out a transcriptional program that includes numerous metabolic enzymes[38], we compared the MYC targets in our dataset of HSP90 metabolic interactome with the universe of MYC targets in the KEGG metabolic pathways. We found a significant overrepresentation of MYC targets in the HSP90 metabolic interactome (Fisher’s exact p < 0.0001, **Fig. 5B**).

**Figure 5.**
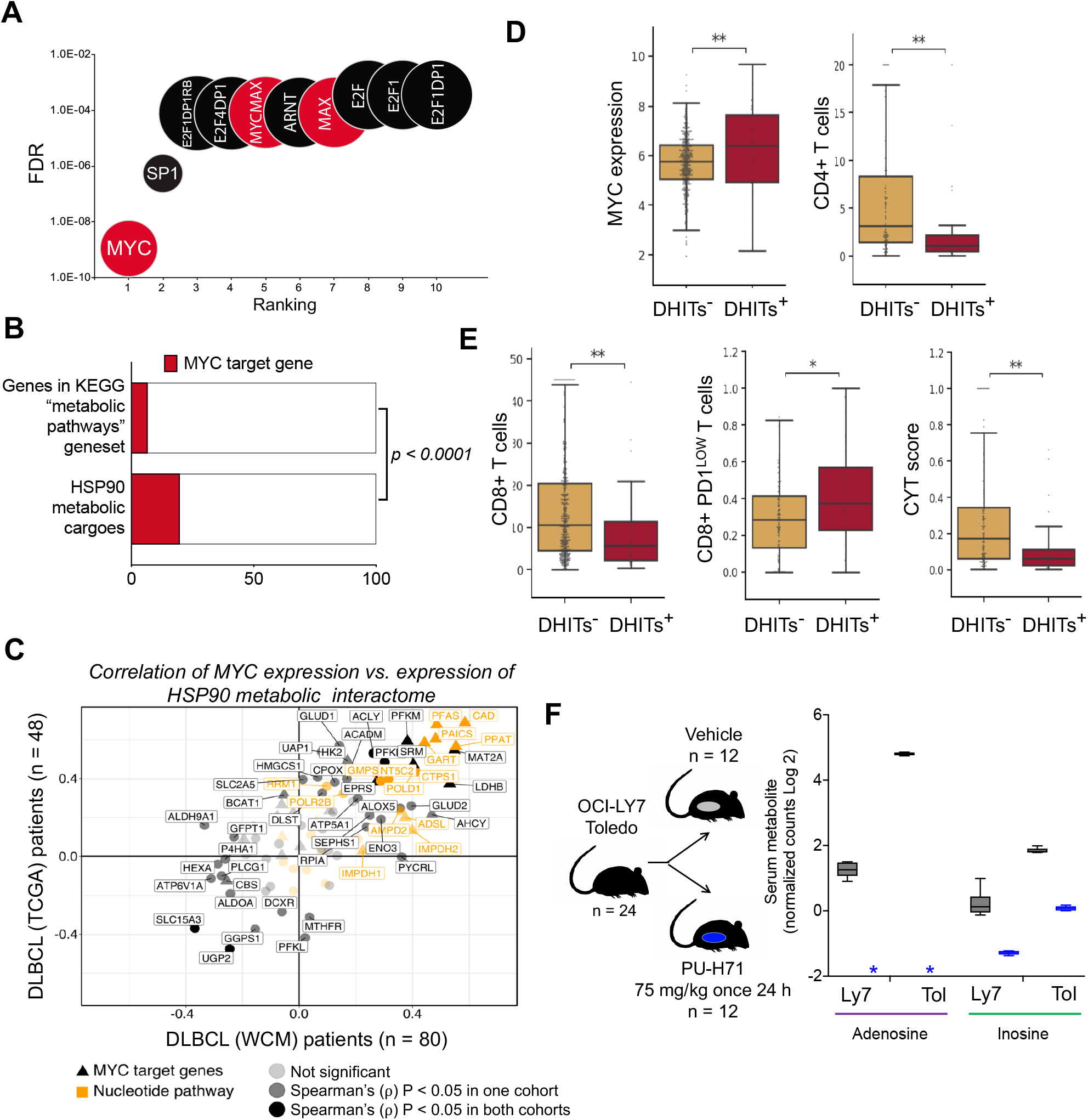
The HSP90 metabolic interactome supports the MYC metabolic program in B-cell lymphomas. **A:** Ranking of transcription factors regulating the HSP90 metabolic interactome in DLBCL according to the presence of canonical binding sites on their promoter regions. Statistical significance was established by FDR. **B:** HSP90 metabolic interactome components presenting MYC binding sites in promoters compared to the universe of metabolic genes from the KEGG database containing MYC binding sites. Statistical enrichment was established by Fisher’s exact test. **C:** Spearman’s rank correlation (ρ) plots comparing the expression of MYC vs. the expression of genes from the HSP90 metabolic interactome. Correlations were conducted in two independent DLBCL patients’ cohorts composed of 48 TCGA cases (Y-axis) and 80 WCM cases (X-axis). Canonical MYC target genes are depicted with triangles. Genes from the nucleotides pathway are shown in yellow. Darker shading indicates significant (p<0.05) correlations found in both cohorts. **D-E:** MYC expression and transcriptomic signature deconvolution of T cells subsets in the lymphoma microenvironment of DLBCL patients with DHIT positive signatures (DHITs+) vs. DHIT negative signatures (DHITs-). **F:** Cartoon representing the mice experiment. Serum metabolite levels of adenosine and inosine in mice treated with PU-H71 or vehicle. P values were calculated by T-test. *<0.05 and **<0.01.

MYC is an important prognosis-associated oncogene in DLBCL[1, 39, 40] that regulates a myriad of cellular pathways. To determine the relevance of the HSP90 metabolic interactome in the metabolic transcriptional program of MYC in DLBCL, we calculated the transcript expression correlations of MYC with the subset of transcripts corresponding to the HSP90 metabolic interactome proteins. We used two independent cohorts of DLBCL patients; our 80 DLBCL RNA-seq samples (WCM cohort) and a publicly-available set of 48 DLBCL RNA-seq samples (TCGA cohort). There were 37 HSP90 metabolic interactome genes that correlated significantly (p-value < 0.05) with MYC expression in both datasets, with 35% of them belonging into the “nucleotides” pathway (**Fig. 5C** and **Table S4**). These included the MYC target genes IMPDH2 of the purine biosynthesis and CTPS1 of the pyrimidine biosynthesis (**Fig. 5C**). Additionally, the correlation between HSP90 metabolic interactome genes and MYC expression was equally robust in 30 DLBCL cell lines (**Fig. S6**).

Recently, a transcriptomic-derived signature identified a group of MYC-driven DLBCLs termed DHITsig positive[40]. Similar to the HSP90 metabolic proteome, DHITsig positive lymphomas are enriched for MYC and E2F target genes (**Fig. 5A**) including metabolic genes associated with oxidative phosphorylation and mTOR signaling[40] as well as lower infiltration of CD4+ T cells. To determine whether DHITsig positive DLBCL could potentially benefit from decreasing the MYC immunometabolic program supported by HSP90, we deconvoluted the T cell signatures from the lymphoma microenvironment. We found that DHITsig positive DLBCL had, as reported, higher expression of MYC and lower infiltration of CD4+ T cells (**Fig. 5D**). In addition, we found a lower infiltration of effector CD8+ T cells in these lymphomas that are characterized by low PD1 expression but low cytotoxicity score (**Fig. 5E**). This profile suggested that a modification of a purine-rich tumor macro and microenvironment could positively impact on the activity of CD8+ T cells[36]. To determine whether HSP90 inhibition decreases the secretion of purines in DHITsig positive lymphomas, we implanted two human DHITsig positive DLBCL cell lines into mice and, once tumor reached 200 mm^3^, we treated them with vehicle or PU-H71 for 24h (**Fig. 5F**). We found that similar to DLBCL patients, low-dose HSP90 inhibitor significantly decreased the serum levels of adenosine and inosine (**Fig. 5F**), suggesting that a more favorable environment for the activity of T cells can be obtained. Our results broadly suggest that that HSP90 inhibition may constitute a potential immune booster in MYC-driven DLBCL.

### MYC activation favors the assembly of metabomes

MYC translocations resulting in high levels of MYC expression are considered a primary genetic event in BL[39]. We thus investigated whether the expression correlation of MYC with the subset of transcripts corresponding to the HSP90 metabolic interactome proteins was also present in BL patients. We analyzed and compared the gene expression profiles of 24 BL patients with 20 primary mediastinal B-cell lymphoma (PMBL) patients, a B-cell lymphoma not driven by MYC[39]. Similar to DLBCL, we found a strong association between the expression of MYC, HSP90 (HSP90AB1) and HSP90 metabolic interactome genes in BL patients, but not in PMBL patients (**Fig. 6A**). Moreover, the nucleotide metabolism enzymes and direct MYC target genes IMPDH2, CTPS1 and CAD were among the most strongly positively correlated with MYC expression in BL patients (**Fig. 6A**). We then analyzed if they were part of HSP90-dependent metabosomes by using PU-H71 bait in sequential purification experiments using primary lymphoma cells obtained from a MYC-translocated BL patient (**Fig. 6B**). CLPP[41], a mitochondrial protein and MYC target gene, was used as negative control whereas HSP90 was used as positive control. We found that, similar to cell lines, in primary BL cells HSP90-dependent metabosomes contain the majority of IMPDH2, CTPS1 and CAD present in the sample (**Fig. 6B**). There was no binding of active HSP90 to CLPP, which reinforces our early observation that PU-H71 is not targeting the mitochondria directly.

**Figure 6.**
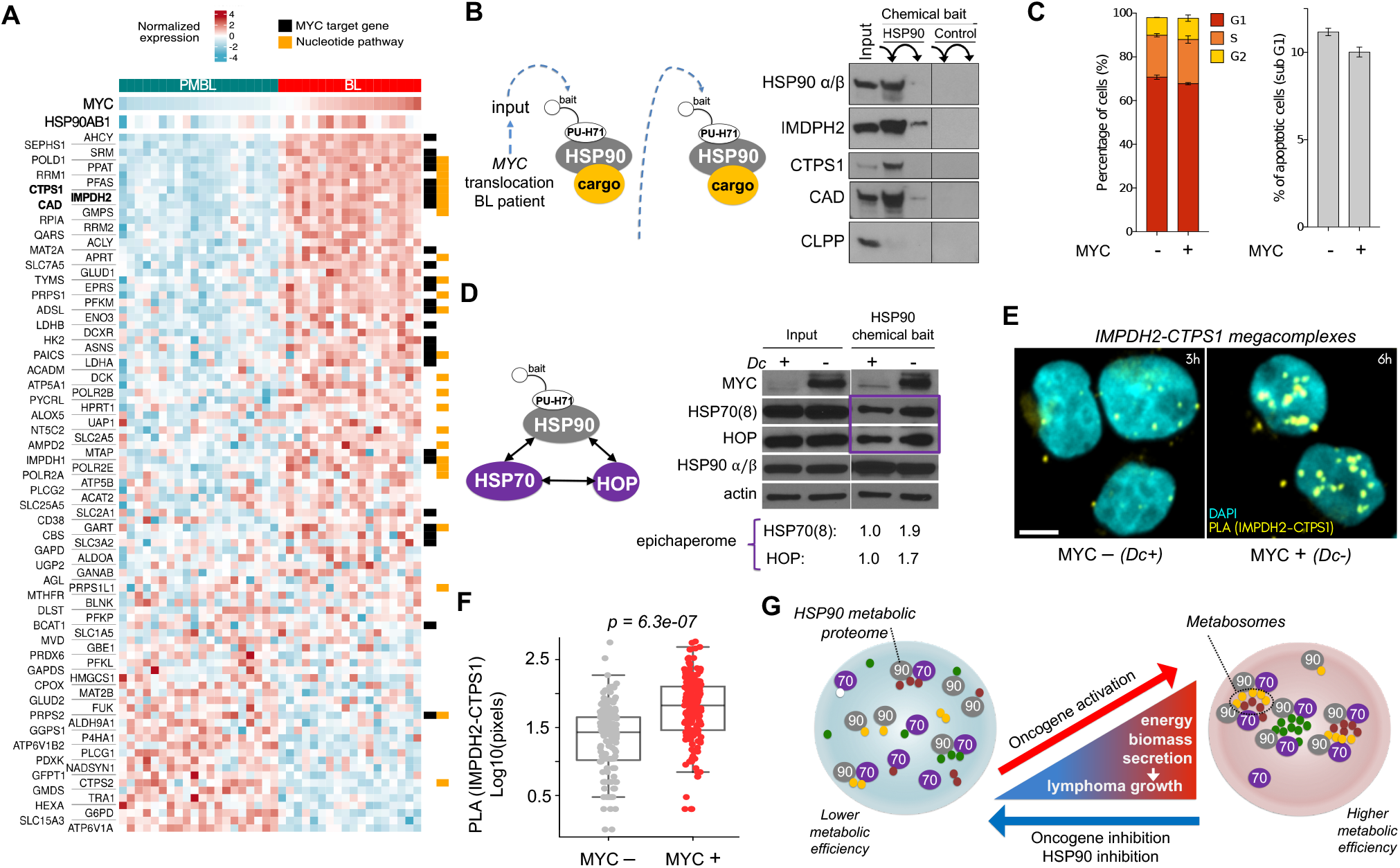
*MYC* activation in lymphoma cells induces metabosome formation and associate with changes in the microenvironment. **A:** Transcript expression heatmap of MYC, HSP90 (HSP90AB1) and HSP90 metabolic interactome components in BL cases (in red) and PMBL cases (in cyan). Canonical MYC target genes are marked with black rectangles and genes from the nucleotides pathway with yellow rectangles. **B:** Validation of HSP90 metabolic interactome components IMPDH2, CTPS1 and CAD in a BL patient sample presenting MYC translocation. HSP90 was used as positive control and CLPP as negative control. Chemically inert beads were used as control beads. **C:** Cell cycle effect and dead cells (sub G1 fraction) by flow-cytometry of P493-6 BL cells in absence of MYC and upon its induction. **D:** Abundance of the epichaperome (interaction of HSP90 and HSP70 chaperomes) in P493-6 BL cells according to MYC induction. Total protein abundance in cells (input) and epichaperome fraction purified with HSP90 chemical beads in P493-6 cells in absence of MYC and upon its induction by doxycycline (Dc) withdrawal (6 h). The quantification at the bottom indicates the abundance of the HSP70- and HOP-containing epichaperome complexes (purple square) normalized to the MYC low state. **E:** Representative imaging of IMPDH2 and CTPS1 endogenous macrocomplexes (yellow puncta) in P493-6 BL cells in absence of MYC and upon its induction by Dc withdrawal (6 h). The bar represents 5 μm. **F:** Quantification of the IMPDH2 and CTPS1 endogenous macrocomplexes in P493-6 BL cells in absence and presence of MYC. **G:** Graphical representation of the findings in this work.

The epichaperome results from changes in the chaperome that are driven by activation of MYC and it manifest as an enhanced physical integration of the HSP90 and HSP70 machineries[27]. To demonstrate the dependence on MYC for epichaperome formation, we compared the epichaperome abundance in B-cell lymphoma cells P493-6 carrying a conditional MYC allele. Epichaperome abundance was determined by co-purification of PU-H71 chemical bait with HSP70 (HSC70) and the adaptor protein HOP (HSC70/HSP90-organizing protein). P493-6 cells were treated with doxycycline (Dc) to downregulate MYC 6 h followed by 1 h Dc withdrawal. In this condition there were no significant changes in cell proliferation and/or apoptosis between MYC- and MYC+ cells (**Fig. 6C**). Whereas the total abundance of HSP90, HSP70 and HOP did not change between MYC+ and MYC-cells, the epichaperome abundance increased in MYC+ cells (**Fig. 6D**); suggesting that MYC activity rapidly increases the nucleating function of HSP90. This translated into changes in the topology of cytosolic metabosomes evidenced by a significant increase in the number of stable IMPDH2-CTPS1 megacomplexes in MYC+ cells (**Fig. 6E-F**). These data indicated that establishment of a MYC-driven metabolism is favored by the presence of HSP90 metabosomes and that HSP90 inhibitors crippled this metabolic program. Taken together, our data demostated that epichaperome inhibition leads to metabosome disassembly causing the ultimate metabolic collapse of lymphoma cells providing an aditional mechanism for the antineoplastic effect of HSP90 inhibitors (**Fig. 6G**).

## DISCUSSION

In a group of cancer subtypes that include DLBCL and BL, HSP90 organizes into higher order heterogeneous chaperome complexes termed epichaperomes that are more stable than the classical chaperomes characteristic of normal cells[27]. Contrary to classical transient chaperomes, whose function is to fold and stabilize proteins, the more permanent epichaperome complexes allow the maintenance of diverse macrocomplexes in active configurations[26, 27]. We report here the discovery of a novel mechanism of metabolic regulation in cancer cells that is characterized by the HSP90-dependent nucleation of enzymatic macrocomplexes. By limiting the cytosolic distribution of functionally-related enzymes, these assemblies (termed metabosomes) increase the efficiency of metabolic pathways. We demonstrate that DLBCL cells with active cytosolic HSP90 possess a metabolic advantage to maintain their biomass and nutrient utilization over DLBCL cells with inhibited HSP90. *MYC* and other transcription factors exert their oncogenic activity by increasing the expression of their target genes. However, there are several limitations to the extent to which a protein can be expressed without inducing detrimental molecular crowding[42]. HSP90 nucleating capacity adds to the regulation of protein crowding by favoring metabosomes thus maximizing the activity of overexpressed enzymes.

Microcompartmentation, either due to membrane-restriction or by clusters of enzymes into functional complexes, has implications for efficiency of metabolic pathways and to segregate “moonlighting” functions of many enzymes and metabolites. In addition to the generation of gradients, the spatial organization of enzymes into complexes has been proposed to facilitate substrate funneling[8]. Metabolism is thus accomplished through the spatial microcompartmentation of metabolic complexes into vesicles and organelles as occurs, for example, during fatty-acid oxidation and the tricarboxylic acid cycle[8]. However, the demonstration of the functional metabolic microcompartmentation of the cytosol in mammalian cells has been more elusive. Cytosolic fluidity is subject to the dense lattice constituted by the cytoskeleton[42] and, accordingly, pharmacological disruption of actin and/or tubulin polymerization decreases megacomplexes formation and cytosolic distribution[14, 43]. However, the dynamic mechanism of organizing cytosolic multi-enzymatic complexes into functional microcompartments, rather than stochastic microcompartments as the mere distribution model imposed by the cytoskeleton would suggest, has remained incompletely understood. Taking advantage of our methodology to isolate native proteins nucleated by active HSP90, we characterized the network of functionally-related metabolic enzymes from the cytosol of DLBCL cells into metabosomes. Similar to other chaperone-containing phase-separated granules[18, 44, 45], it is possible that the transient, enzyme-enzyme interactions during catalysis become more stable in the context of HSP90-containing epichaperomes[26, 27].

There is now intense interest in targeting aberrant tumor cell metabolism for cancer therapeutics. However, with the possible exception of selective compounds targeting mutant enzymes, drugs that can safely target metabolic pathways in tumors, as opposed to normal tissues, have not been clearly identified[1]. Our study supports the notion of targeting not a single protein but, rather, metabosomes to effectively decrease tumor metabolism. Overall decrease in tumor metabolism with HSP90 inhibitors is achieved earlier and at lower doses than those required to elicit protein degradation and apoptosis[29], a hallmark of all HSP90 inhibitors. This will offer the advantage of lower toxicity to normal tissues since normal cells do not contain epichaperomes[26, 27], and could also decrease the emergence of cell-intrinsic compensatory pathways. Moreover, this approach could even induce a cancer cell metabolic reprogramming that in turn could improve the anti-neoplastic activity of other therapeutic approaches targeting lymphoma cells or directed to the lymphoma immune environment. In this regard, our study shows that HSP90 activity in DLBCLs favors an inosine-rich environment, that we found prevalent in DLBCL patients. Inosine is an endogenous nucleoside produced by deamination of adenosine by ADA. Although inosine has a prolonged half-life in comparison with adenosine, neither nucleoside is usually detected in the plasma of healthy individuals. Through the direct activation of purinergic receptors in immune cells, inosine exerts *in vitro* and *in vivo* immunomodulatory effects similar to those described for adenosine[46]. Therefore, our results propose that HSP90 inhibitors administered at relatively low-dose could deplete the lymphoma environment from the immunosuppressive effects of inosine and thus potentially improve the lymphoma immune response.

We also showed that the regulation of cellular metabolism through this mechanism is particularly relevant for lymphomas with MYC activation such as aggressive sub-types of lymphomas such as DHITsig positive DLBCL and BL. The activity of HSP90 is required to support the metabolic reprogramming driven by MYC/MAX. In cancer cells, MYC enhances both catabolism (i.e., the breakdown of nutrients to produce ATP and reductive power such as NADPH) and anabolism (i.e., the uptake of nutrients to biosynthesize macromolecules) to meet the challenges of tumor growth and proliferation[38]. We have demonstrated that HSP90 is required to optimize several cytosolic MYC metabolic pathways, and in particular the production of nucleotides. Other MYC activated pathways such as fatty acids and cholesterol metabolism seem to be less dependent on the nucleating properties of HSP90 in B-cell lymphomas, probably because they take place at membranes or membrane-compartmentalized organelles. Another relevant factor is that unlike fatty acids and amino acids, nucleotides may not be provided in sufficient amounts or proportions by the microenvironment, and their synthesis requires a combination of non-essential amino acids, ribose and one-carbon donors, all of which are affected by HSP90 inhibition to a greater or lesser extent.

In sum, our work demonstrates that the HSP90 acts as multimolecular scaffold in B-cell lymphomas, providing cytosolic components of the metabolism with a framework on which they can work more efficiently than they would without HSP90 participation. This function was therapeutically exploited by using PU-H71, an inhibitor preferentially targeting HSP90 contained in epichaperomes, to depress the tumor metabolism affecting biomass gain and secretory functions. Moreover, in MYC-driven lymphomas this strategy could potentially be used to improve the activity of immune cell-targeting approaches. Given the importance of MYC in driving metabolic reprogramming in other cancer types, our results bear promises as an alternative manner to target the aberrant metabolism characteristic of cancer cells.

## MATERIALS AND METHODS

### Cell lines studies

Lymphoma cell lines OCI-LY1 and OCI-LY7 were cultured in Iscove’s modified Eagle Medium (IMDM) with 20% fetal bovine serum (FBS). Karpas422, Toledo, SU-DHL-6 and Raji were cultured in RPMI-1640 with 10% FBS and 1% HEPES. These cell lines were obtained from the Ontario Cancer Institute, DMSZ or ATCC and regularly tested for *Mycoplasma sp*. contamination by PCR. Annual cell identification was carried out by single nucleotide polymorphism. P493-6 cells were obtained from the original developer Dr. Dirk Eick. Cells were maintained with 1% penicillin/streptomycin in a 37 °C, 5% CO2, humidified incubator.

### Mouse studies

All animal procedures were approved by the Research Animal Resource Center of WCM. OCI-LY7 and Toledo DLBCL cells were injected in the flank of 24 severe combined immunodeficiency (SCID) mice purchased from the US National Cancer Institute (NCI). Once tumors reached 200 mm^3^, mice were treated with vehicle (0.9% sodium chloride solution, n = 6 for each cell line) or 75 mg/kg PU-H71 (n = 6 for each cell line) for 24 h. At the end of the experiment, mice were euthanized by CO2 inhalation and blood obtained post-mortem from cardiac puncture.

### Human studies

Serum from de-identified healthy volunteers (n = 25) and DLBCL patients (n = 50) were obtained with the approvals of the IRBs of Weill Cornell Medicine (New York, USA) and Saint Louis University Hospital (Paris, France). Serum from healthy volunteers were selected to matched the gender and age distribution of DLBCL patients in a 1:2 ratio. Sera were kept at −80 C till the moment of processing for metabolomics. De-identified BL tissue (n = 1) and DLBCL tissues (n = 80) were obtained with the approvals of the IRBs of Weill Cornell Medicine (New York, USA) and University of Turin (Turin, Italy). Tissues were kept at −140C till processing. Plasma from DLBCL patients enrolled in the clinical trial NCT01393509 were obtained with the approval of the IRB of Memorial Sloan Kettering Cancer Center (New York, USA).

### Cell number determination

All cell lines were grown at densities that allowed to maintain vehicle-treated cells in exponential growth over the drug exposure time. Relative cell number was determined by a fluorescence assay based on the measurement of a constitutive protease activity restricted to intact viable cells and that is independent of the cellular metabolic activity (CellTiter-Fluor, Promega). The fluorescence signal was kept within the linear range and was proportional to cell number. For the measurements we used a Synergy4 microplate reader (BioTek).

### Chemical affinity purification of the HSP90 interactome

PU-H71 and control (2-methoxyethlamine) beads were synthesized as previously reported[47]. Before use, beads were washed 3 times in Felts buffer (20 mM HEPES, 50 mM KCl, 5 mM MgCl2, 0.01% NP-40, freshly prepared 20 mM Na_2_MoO_4_, and cOmplete protease inhibitors; Roche). 10-20 million cells were lysed in 0.5 mL Felts buffer by rotation at 4°C for 30 minutes and vortexing for 10 seconds, followed by centrifugation at 14,000 g for 5 minutes at 4°C. Supernatants were precleared with 100 μL of control beads for 1 hour at 4°C. After preclearing, half of each sample was incubated with 100 μl of either control or PU-H71 beads at 4°C overnight in the dark. Following incubation, bead conjugates were washed 3 times with Felts buffer, boiled in Laemmli buffer and resolved by SDS-PAGE by standard immunoblotting procedure. The proteomics methodology was published in[30].

### Immunoblotting

Cells were lysed in protease inhibitor-supplemented RIPA buffer by rotation at 4°C for 30 minutes, followed by centrifugation at 14,000 g for 5 minutes at 4°C. Protein concentration was determined by BCA assay (Pierce Biotechnology) according to manufacturer’s instructions. Protein lysates (10 - 20 μg) were resolved by SDS-PAGE, transferred to PVDF (polyvinylidene difluoride) membranes, and probed with the indicated primary antibodies: α-tubulin (ab4074, Abcam), β-actin (AC-15, A3854, Sigma), BLNK (2B11, sc-8003, Santa Cruz), CAD (RB18384, AP11110c-ev, Abgent), CLPP (B-12, sc-271284, Santa Cruz), CTPS1 (C-13, sc-131474, Santa Cruz), G6PD (ab76598, Abcam), GAPDH (6C5, MAB374, Millipore Sigma), Hexokinase 2 (C-14, sc-6521, Santa Cruz), HSP70 (W27, sc-24, Santa Cruz), HSP90 α/β (F-8, sc-13119, Santa Cruz), IMPDH2 (EPR8365B, ab129165, Abcam), and MTAP (EPR6893, ab126770, Abcam). Membranes were then incubated with peroxidase conjugated secondary antibodies (sc-2004 and sc-2005, Santa Cruz). We used ECL Western Blotting Substrate (Pierce Biotechnology) according to manufacturer’s instructions and the blots were visualized by autoradiography. Quantitative densitometry analysis of western blot bands was performed employing Image J version 10.2 (NIH).

### Metabolomic assays

*Cells:* Low-passage OCI-LY1 and OCI-LY7 cell cultures were maintained at 10^6^ cells/mL in fresh medium the night before adding either vehicle (PBS) or PU-H71 (0.5 μM) to the culture for 6 hours. At endpoint, cells were harvested, washed once with cold PBS and snap frozen in nitrogen. Untargeted metabolomics was run by Metabolon Inc. Briefly, measurements were performed by ultra-high-performance liquid-phase chromatography and gas-chromatography separation, coupled with tandem mass spectrometry. For validation, targeted profiling of polar metabolites was performed at Weill Cornell Medicine Proteomics and Metabolomics Core Facility (WCM-PMCF) by ultra-high-performance liquid-phase chromatography coupled with tandem mass spectrometry followed by metabolite identification and quantitation. For the overlaying of proteins and metabolites onto the KEGG map of human metabolism we used iPath[48]. *Mice:* total blood was obtained post-mortem from experimental mice by cardiac puncture and immediately processed to isolate the serum fraction. Serum was processed for nucleotide analysis in the WCM-PMCF as before.

### Protein proximity ligation assay (PLA)

P493-6 cells in absence of MYC and upon its induction by doxycycline withdrawal (6 h) and OCI-LY1 cells treated with vehicle or 500 nM PU-H71 for 12 h, were spun down at 250 g for 5 minutes, plated on 8 mm coverslips previously coated with Cell-Tak and incubated for 15 minutes at 37 °C to adhere. After washing with PBS, they were fixed with 4 % paraformaldehyde and permeabilized with PBS-Tween 0.5%. Cells were then subjected to the PLA assay using the Duolink red kit (O-link Bioscience, Sweden) in a humidity chamber according to the manufacturer’s instructions. Briefly, cells were blocked with Duolink blocking buffer for 30 m and incubated for 1 h at 37 °C in the presence of primary antibodies for IMPDH (F-6, sc-166551, Santa Cruz) and CTPS1 (sc-131474, Santa Cruz) diluted in Duolink antibody diluent at 1:50 and 1:100, respectively. After washing with Duolink washing buffer, cells were incubated with plus and minus PLA probes for 1 h at 37 °C. After washing with Duolink washing buffer, ligase was added to the samples for 30 minutes at 37 °C. After washing, samples were then incubated in the presence of Duolink polymerase buffer containing the polymerase for 100 minutes at 37 °C. Finally, covers were washed and mounted with Fluoromount-G (Electron Microscopy Sciences) for laser confocal confocal microscopy. Cell images were acquired using a confocal microscope (Zeiss LSM880) with a Plan-Apochromat 63x/1.40 oil objective. In order to scan the whole cellular volume, 10 images at 0.2-micron step size were taken per field. Only clearly identifiable dots in the maximum intensity Z projection within the cell boundary were considered as indicators of protein proximity.

### Mass accumulation rate

We measured the mass accumulation rates (MAR) of individual cells using a serial suspended microchannel resonator (sSMR) device, as described previously[35, 49]. OCI-LY1 cells were cultured in IMDM with 20% FBS, then resuspended in their own medium at a concentration of 0.5E6 cells/mL for measurement. Cells were sampled from this untreated population and weighed 12 times over a 30-minute interval as they flowed through the sSMR device. After 1.5 hours, 1 μM PU-H71 was added to the cell suspension and sampling of the population by the sSMR continued for ~4 hours. A total of 106 cells were measured pre-treatment and 161 cells post-treatment.

### Substrates consumption and lactate secretion

Low-passage cells were washed twice in PBS and resuspended at 10^6^ cells/mL in RPMI without glucose or glutamine supplemented with 5.55 mM glucose, 1 mM glutamine and 1 % HEPES. A well without cells and with the same medium under the same culture conditions was used as reference to estimate the glucose and glutamine consumed and the lactate secreted in the wells with cells. For the measurements we centrifuged the cells at 250 g, 4 °C, for 5 minutes and immediately snap froze the supernatant to avoid degradation until analyzing it in a BioProfile Basic (Nova Biomedical) according to manufacturer’s instructions.

### Oxygen consumption rate (OCR) and extracellular acidification rate (ECAR)

OCR and ECAR were measured using the Seahorse XF96 Extracellular Flux Analyzer (Agilent). Low-passage cells were seeded on Cell-Tak (BD Biosciences) pre-coated XF96 plates at 10^5^ cells/well in 175 μL of sodium bicarbonate-free XF Base (Agilent) media freshly supplemented with 10 mM glucose, 2 mM L-glutamine and 1 mM pyruvate. Plates were spun at 40 g with breaks off and incubated in the XF incubator without CO^2^ for 25 min to ensure cell attachment. Measurements were taken before the addition of any inhibitor (basal respiration) and after the sequential injection of 1 μM oligomycin (for ATP-linked respiration), 0.5 μM FCCP (for maximal respiration) and 1 μM rotenone/antimycin A (for non-mitochondrial oxygen consumption).

### Medium oxygen consumption

Medium oxygen levels were measured in an SDR SensorDish Reader (PreSens). Cells were washed twice with PBS and changed to RPMI without glucose and glutamine (Biological Industries) and with 10% dialyzed FBS (Gibco) at 10^6^ cells/mL. They were plated at 0.8 mL of culture/well on pre-warmed 24-well OxoDish (PreSens) plates with integrated oxygen sensors. Medium oxygen levels were measured for 40 minutes to control for inter-well variability. The measurements were then paused to add either vehicle or 0.5 μM PU-H71 to the corresponding wells, and the oxygen of the medium was monitored for 1 more hour, at which point the measurement was paused again to add either PBS or 2 mM glutamine (Gibco) to the corresponding wells. Measurements went on for up to a total of 4 hours under regular incubation conditions (37 °C, 5% CO2). The measurements after addition of substrate were normalized to 1 by dividing by the levels of each well before addition of substrate.

### DNA and protein syntheses

Low-passage cell cultures were maintained at 10^6^ cells/mL in fresh medium and either vehicle (PBS) or PU-H71 were added to the culture for 5 hours, after which the thymidine analog EdU was added (10 μM). After 1 hour, 10^6^ cells were harvested and processed according to manufacturer’s instructions. DAPI was added (1 μg/mL) to the final cell suspension and cells were analyzed after at least overnight incubation for best results. The DNA synthesis data is the mean fluorescence of the EdU incorporated in S phase cells. For protein synthesis, cells were treated similarly with vehicle or PU-H71 (5.5 hours). Then they were changed to RPMI without methionine (Gibco) and supplemented with 50 μM of the methionine analog HPG for 30 minutes. They were harvested and processed according to manufacturer’s instructions. The protein synthesis data is the mean fluorescence of the HPG incorporated in the healthy population of cells. In both cases, data were collected on a MACSQuant flow cytometer (Miltenyi Biotec) and analyzed using FlowJo version 10.0.6 (Tree Star Inc.).

### Real-time reverse transcriptase-qPCR

Total RNA was purified using TRIzol Reagent (Thermo Fisher Scientific) following manufacturer’s instructions and resuspended in RNAse-free water. cDNA was synthesized using high capacity RNA-to-cDNA kit (Applied Biosystems). SYBR Green FastMix (Quanta BioSciences) was used for the PCRs. Primer sequences were designed to span exon-exon junctions in Primer-BLAST (NCBI).

### Metabolic tracing

OCI-Ly1 cells were maintained in Iscove’s Modified Dulbecco’s Medium (IMDM) supplemented with 10% FBS and 1 % Penicillin/Streptomycin until a density of 1 x 10^6^ cells per mL was reached. Cells were then washed once with PBS and incubated in glucose-depleted Dulbecco’s Modified Eagle Medium (DMEM) for 1 h prior to starting the tracing. After washing once with PBS, cells were incubated for 30 min, 3 h or 6 h in the presence of either DMSO (vehicle) or PU-H71 (1 μM) in conditional DMEM containing 2.5 mM of tracer U-13 C6-glucose plus 10% dialyzed FBS. After the incubation time, cells were washed once with PBS and centrifuged, and the pellets were saved at −80°C. Metabolites were extracted from cells using 80% methanol. Targeted LC/MS analyses were performed on a Q Exactive Orbitrap mass spectrometer (Thermo Scientific) coupled to a Vanquish UPLC system (Thermo Scientific). The Q Exactive operated in polarity-switching mode. A Sequant ZIC-HILIC column (2.1 mm i.d. × 150 mm, Merck) was used for separation of metabolites. Flow rate was 150 μL/min. Buffers consisted of 100% acetonitrile for A, and 0.1% NH4OH/20 mM CH3COONH4 in water for B. Gradient ran from 85 to 30% A in 20 min followed by a wash with 30% A and reequilibration at 85% A. Metabolites were identified on the basis of exact mass within 5 ppm and standard retention times. Metabolites and their 13C isotopologues were identified on the basis of standard retention times and exact mass within 5 ppm. Relative quantitation was performed based on metabolite peak area. The studies were conducted by the WCM Metabolomics and Proteomics Facility and data analysis was done using in-house written scripts.

### Isolation of human B cell populations

For isolation of primary human B cell populations, we used de-identified human tonsillectomy specimens from New York Presbyterian Hospital with approval from Weill Cornell Medicine Institutional Review Board. Tonsils were minced, and mononuclear cells were isolated using Ficoll Histopaque density centrifugation. Naïve B cells were separated by positive selection using AutoMACS system (Miltenyi Biotec) after incubation with anti-IgD-FITC (BD Pharmingen) followed by anti-FITC microbeads (Miltenyi Biotec). Germinal center B cells (GCB) were separated by positive selection with anti-CD77 (AbD Serotec) followed by mouse anti-IgM, IgG1 isotype (BD Pharmingen) and anti-mouse-IgG1 microbeads (Miltenyi Biotec). Naive and germinal center B cell purity (>90%) was determined by flow cytometry analysis of surface IgD (BD Biosciences 555778), CD77 (BioRad MCA579) and CD38 (BD Biosciences 340439). GCB were further verified by expression of A4GALT and BCL6 by RT-qPCR.

### Statistical analysis

#### Metabolomics dataset analyses

All metabolite concentrations were converted to log2 prior to statistical analysis, since previous studies have demonstrated that metabolites are largely log-normally distributed. Identified metabolites with more than 20% missing values were excluded. The remaining missing values were imputed by drawing from a normal distribution with the same mean and standard deviation as the non-missing values from the respective metabolite. No named metabolites and exogenous metabolites were excluded from analysis. Each metabolite was annotated with a) one of seven major biochemical super-pathways (“amino acid”, “peptide”, “lipid”, “energy”, “carbohydrate”, “nucleotide” and “cofactors & vitamins”), and b) one sub-pathway. For each group of metabolites belonging to a pathway, mean z-scores were computed as a measure of average activity over all members of the pathway. Differences in metabolites between experimental conditions were assessed by standard two-sample t-tests.

#### Transcriptional datasets analyses

For the TCGA DLBCL set, level 3 raw expression counts of 48 patients were downloaded from TCGA. The WCM DLBCL cohort and cell lines RNA-seq data were aligned using STAR (v2.3) to human reference genome (version hg19/GRCh37), and raw counts were calculated with HT-seq. Normalized read counts as provided by DESeq2 were extracted from the three datasets for posterior analyses. Spearman correlations were calculated in R and plotted with ggplot2. 44 samples corresponding to BL and PMBL profiled on Affymetrix HG U133 Plus 2 arrays were downloaded from http://llmpp.nih.gov/BL/ and were used as deposited by the authors.

#### Proteomics dataset analyses

Data from the chemical precipitation of HSP90 cargoes from the cytoplasm of DLBCL cell lines OCI-LY1 and OCI-LY7 was obtained from previous experiments in our lab[30] and the union of the HSP90 interactome in the two cell lines was analyzed by KEGG pathway enrichment analysis in WebGestalt[50]. The HSP90 cargo proteins under the category “metabolism” were further analyzed by STRING performed with a minimum required confidence interaction score of 0.4 on a scale from 0 to 1, and by determining the top 10 overrepresented KEGG pathways.

#### Degree of association between proteins and metabolites

To compare sets of significantly changed metabolites and the HSP90 metabolic interactome, metabolites and proteins were mapped to the Recon 2.04 metabolic network. This curated network currently contains 5,063 metabolites, involved in 7,440 metabolic reactions, annotated with 2,140 genes. Each reaction is annotated with one or more metabolic pathways such as “pentose phosphate” and “fatty acid oxidation”. In our analysis, all pathway annotations involving transport reactions (e.g. “transport, extracellular”) as well as the generic pathway “exchange/demand reaction” were ignored, since they group metabolites in a generic fashion that does not emphasize actual cellular function. A metabolite and a protein were considered to be “connected” if their respective annotated reaction occurred in the same RECON2 pathway. Connections between metabolites and proteins were visualized in matrix form and as a network.

#### Protein structural and biochemical properties analysis

We used ProtParam to compute the following protein properties: instability index, aliphatic index and number of amino acids. We used UniProtKB to retrieve the adenosine nucleotide binding information. And we used PASTA2 to compile secondary structure predictions and intrinsic disorder.

### Data and software availability

RNA-sequencing data for WCM DLBCL patients (N = 80) and DLBCL cell lines (N = 30) are available at Gene Expression Omnibus (GEO) with the accession number: GSE145043. The metabolomics dataset of DLBCL patients and healthy volunteers is available as Table S3.

## ACKNOWLEDGEMENTS

We thank Dr. Dirk Eick from the GSF-Research Center for Environment and Health, Munich, Germany, for providing P493-6 cells and Drs. Justin Cross and Vladimir Yong from the Marron Cancer Metabolism Center of Memorial Sloan-Kettering Cancer Center for help with cellular OCR and ECAR measurements. The work involving human specimens was supported by the Doris Duke Charitable Foundation DDCF 2012070 (to L.C.). Additional work was supported by the Leukemia & Lymphoma Society SCOR award (LLS-SCOR 7012-16) component core C (to L.C.), the National Institutes of Health 1U54CA217377 (to S.R.M.) and the National Institutes of Health - National Cancer Institute R01CA242069 (to L.C.).

## REFERENCES

1. Calvo-Vidal MN, Cerchietti L. The metabolism of lymphomas. Curr Opin Hematol. 2013;20(4):345–54. Epub 2013/05/16. doi: 10.1097/MOH.0b013e3283623d16. PubMed PMID: 23673340.

2. Caro P, Kishan AU, Norberg E, Stanley IA, Chapuy B, Ficarro SB, et al. Metabolic signatures uncover distinct targets in molecular subsets of diffuse large B cell lymphoma. Cancer Cell. 2012;22(4):547–60. Epub 2012/10/20. doi: 10.1016/j.ccr.2012.08.014. PubMed PMID: 23079663; PubMed Central PMCID: PMCPMC3479446.

3. Vajpayee N, Thakral C, Gopaluni S, Newman N, Gajra A. Activation of mammalian target of rapamycin in diffuse large B-cell lymphoma: a clinicopathological study. Leuk Res. 2012;36(11):1403–9. Epub 2012/08/21. doi: 10.1016/j.leukres.2012.07.016. PubMed PMID: 22902049.

4. Dang CV, Lewis BC. Role of Oncogenic Transcription Factor c-Myc in Cell Cycle Regulation, Apoptosis and Metabolism. J Biomed Sci. 1997;4(6):269–78. Epub 2002/10/19. PubMed PMID: 12386373.

5. Sander S, Calado DP, Srinivasan L, Kochert K, Zhang B, Rosolowski M, et al. Synergy between PI3K signaling and MYC in Burkitt lymphomagenesis. Cancer Cell. 2012;22(2):167–79. Epub 2012/08/18. doi: 10.1016/j.ccr.2012.06.012. PubMed PMID: 22897848; PubMed Central PMCID: PMCPMC3432451.

6. Boroughs LK, DeBerardinis RJ. Metabolic pathways promoting cancer cell survival and growth. Nat Cell Biol. 2015;17(4):351–9. Epub 2015/03/17. doi: 10.1038/ncb3124. PubMed PMID: 25774832; PubMed Central PMCID: PMCPMC4939711.

7. O’Connell JD, Zhao A, Ellington AD, Marcotte EM. Dynamic reorganization of metabolic enzymes into intracellular bodies. Annu Rev Cell Dev Biol. 2012;28:89–111. Epub 2012/10/13. doi: 10.1146/annurev-cellbio-101011-155841. PubMed PMID: 23057741; PubMed Central PMCID: PMCPMC4089986.

8. Schmitt DL, An S. Spatial Organization of Metabolic Enzyme Complexes in Cells. Biochemistry. 2017;56(25):3184–96. Epub 2017/06/06. doi: 10.1021/acs.biochem.7b00249. PubMed PMID: 28580779; PubMed Central PMCID: PMCPMC5574030.

9. Meyer P, Cecchi G, Stolovitzky G. Spatial localization of the first and last enzymes effectively connects active metabolic pathways in bacteria. BMC Syst Biol. 2014;8:131. Epub 2014/12/17. doi: 10.1186/s12918-014-0131-1. PubMed PMID: 25495800; PubMed Central PMCID: PMCPMC4279816.

10. Dhar A, Samiotakis A, Ebbinghaus S, Nienhaus L, Homouz D, Gruebele M, et al. Structure, function, and folding of phosphoglycerate kinase are strongly perturbed by macromolecular crowding. Proc Natl Acad Sci U S A. 2010;107(41):17586–91. Epub 2010/10/06. doi: 10.1073/pnas.1006760107. PubMed PMID: 20921368; PubMed Central PMCID: PMCPMC2955104.

11. Hu H, Juvekar A, Lyssiotis CA, Lien EC, Albeck JG, Oh D, et al. Phosphoinositide 3-Kinase Regulates Glycolysis through Mobilization of Aldolase from the Actin Cytoskeleton. Cell. 2016;164(3):433–46. Epub 2016/01/30. doi: 10.1016/j.cell.2015.12.042. PubMed PMID: 26824656; PubMed Central PMCID: PMCPMC4898774.

12. Shearwin K, Nanhua C, Masters C. Interactions between glycolytic enzymes and cytoskeletal structure--the influence of ionic strength and molecular crowding. Biochem Int. 1990;21(1):53–60. Epub 1990/01/01. PubMed PMID: 2143653.

13. Volker KW, Reinitz CA, Knull HR. Glycolytic enzymes and assembly of microtubule networks. Comp Biochem Physiol B Biochem Mol Biol. 1995;112(3):503–14. Epub 1995/11/01. PubMed PMID: 8529027.

14. An S, Deng Y, Tomsho JW, Kyoung M, Benkovic SJ. Microtubule-assisted mechanism for functional metabolic macromolecular complex formation. Proc Natl Acad Sci U S A. 2010;107(29):12872–6. Epub 2010/07/10. doi: 10.1073/pnas.1008451107. PubMed PMID: 20615962; PubMed Central PMCID: PMCPMC2919939.

15. Castellana M, Wilson MZ, Xu Y, Joshi P, Cristea IM, Rabinowitz JD, et al. Enzyme clustering accelerates processing of intermediates through metabolic channeling. Nat Biotechnol. 2014;32(10):1011–8. Epub 2014/09/30. doi: 10.1038/nbt.3018. PubMed PMID: 25262299; PubMed Central PMCID: PMCPMC4666537.

16. Amar P, Legent G, Thellier M, Ripoll C, Bernot G, Nystrom T, et al. A stochastic automaton shows how enzyme assemblies may contribute to metabolic efficiency. BMC Syst Biol. 2008;2:27. Epub 2008/03/28. doi: 10.1186/1752-0509-2-27. PubMed PMID: 18366733; PubMed Central PMCID: PMCPMC2322945.

17. Srere PA. Complexes of sequential metabolic enzymes. Annu Rev Biochem. 1987;56:89–124. Epub 1987/01/01. doi: 10.1146/annurev.bi.56.070187.000513. PubMed PMID: 2441660.

18. Jin M, Fuller GG, Han T, Yao Y, Alessi AF, Freeberg MA, et al. Glycolytic Enzymes Coalesce in G Bodies under Hypoxic Stress. Cell Rep. 2017;20(4):895–908. Epub 2017/07/27. doi: 10.1016/j.celrep.2017.06.082. PubMed PMID: 28746874; PubMed Central PMCID: PMCPMC5586494.

19. Jeon M, Kang HW, An S. A Mathematical Model for Enzyme Clustering in Glucose Metabolism. Sci Rep. 2018;8(1):2696. Epub 2018/02/11. doi: 10.1038/s41598-018-20348-7. PubMed PMID: 29426820; PubMed Central PMCID: PMCPMC5807315.

20. An S, Kumar R, Sheets ED, Benkovic SJ. Reversible compartmentalization of de novo purine biosynthetic complexes in living cells. Science. 2008;320(5872):103–6. Epub 2008/04/05. doi: 10.1126/science.1152241. PubMed PMID: 18388293.

21. Zhao H, Chiaro CR, Zhang L, Smith PB, Chan CY, Pedley AM, et al. Quantitative analysis of purine nucleotides indicates that purinosomes increase de novo purine biosynthesis. J Biol Chem. 2015;290(11):6705–13. Epub 2015/01/22. doi: 10.1074/jbc.M114.628701. PubMed PMID: 25605736; PubMed Central PMCID: PMCPMC4358094.

22. French JB, Jones SA, Deng H, Pedley AM, Kim D, Chan CY, et al. Spatial colocalization and functional link of purinosomes with mitochondria. Science. 2016;351(6274):733–7. Epub 2016/02/26. doi: 10.1126/science.aac6054. PubMed PMID: 26912862; PubMed Central PMCID: PMCPMC4881839.

23. Carcamo WC, Calise SJ, von Muhlen CA, Satoh M, Chan EK. Molecular cell biology and immunobiology of mammalian rod/ring structures. Int Rev Cell Mol Biol. 2014;308:35–74. Epub 2014/01/15. doi: 10.1016/B978-0-12-800097-7.00002-6. PubMed PMID: 24411169.

24. Pedley AM, Karras GI, Zhang X, Lindquist S, Benkovic SJ. Role of HSP90 in the Regulation of de Novo Purine Biosynthesis. Biochemistry. 2018;57(23):3217–21. Epub 2018/03/20. doi: 10.1021/acs.biochem.8b00140. PubMed PMID: 29553718.

25. French JB, Zhao H, An S, Niessen S, Deng Y, Cravatt BF, et al. Hsp70/Hsp90 chaperone machinery is involved in the assembly of the purinosome. Proc Natl Acad Sci U S A. 2013;110(7):2528–33. Epub 2013/01/30. doi: 10.1073/pnas.1300173110. PubMed PMID: 23359685; PubMed Central PMCID: PMCPMC3574928.

26. Joshi S, Wang T, Araujo TLS, Sharma S, Brodsky JL, Chiosis G. Adapting to stress-chaperome networks in cancer. Nat Rev Cancer. 2018. Epub 2018/05/26. doi: 10.1038/s41568-018-0020-9. PubMed PMID: 29795326.

27. Rodina A, Wang T, Yan P, Gomes ED, Dunphy MP, Pillarsetty N, et al. The epichaperome is an integrated chaperome network that facilitates tumour survival. Nature. 2016;538(7625):397–401. Epub 2016/10/21. doi: 10.1038/nature19807. PubMed PMID: 27706135; PubMed Central PMCID: PMCPMC5283383.

28. Moulick K, Ahn JH, Zong H, Rodina A, Cerchietti L, Gomes DaGama EM, et al. Affinity-based proteomics reveal cancer-specific networks coordinated by Hsp90. Nat Chem Biol. 2011;7(11):818–26. Epub 2011/09/29. doi: 10.1038/nchembio.670. PubMed PMID: 21946277; PubMed Central PMCID: PMCPMC3265389.

29. Cerchietti LC, Lopes EC, Yang SN, Hatzi K, Bunting KL, Tsikitas LA, et al. A purine scaffold Hsp90 inhibitor destabilizes BCL-6 and has specific antitumor activity in BCL-6-dependent B cell lymphomas. Nat Med. 2009;15(12):1369–76. Epub 2009/12/08. doi: 10.1038/nm.2059. PubMed PMID: 19966776; PubMed Central PMCID: PMCPMC2805915.

30. Goldstein RL, Yang SN, Taldone T, Chang B, Gerecitano J, Elenitoba-Johnson K, et al. Pharmacoproteomics identifies combinatorial therapy targets for diffuse large B cell lymphoma. J Clin Invest. 2015;125(12):4559–71. Epub 2015/11/04. doi: 10.1172/JCI80714. PubMed PMID: 26529251; PubMed Central PMCID: PMCPMC4665772.

31. Culjkovic-Kraljacic B, Fernando TM, Marullo R, Calvo-Vidal N, Verma A, Yang S, et al. Combinatorial targeting of nuclear export and translation of RNA inhibits aggressive B-cell lymphomas. Blood. 2016;127(7):858–68. Epub 2015/11/26. doi: 10.1182/blood-2015-05-645069. PubMed PMID: 26603836; PubMed Central PMCID: PMCPMC4760090.

32. Leuenberger P, Ganscha S, Kahraman A, Cappelletti V, Boersema PJ, von Mering C, et al. Cell-wide analysis of protein thermal unfolding reveals determinants of thermostability. Science. 2017;355(6327). Epub 2017/02/25. doi: 10.1126/science.aai7825. PubMed PMID: 28232526.

33. Thiele I, Swainston N, Fleming RM, Hoppe A, Sahoo S, Aurich MK, et al. A community-driven global reconstruction of human metabolism. Nat Biotechnol. 2013;31(5):419–25. Epub 2013/03/05. doi: 10.1038/nbt.2488. PubMed PMID: 23455439; PubMed Central PMCID: PMCPMC3856361.

34. Taldone T, Patel PD, Patel M, Patel HJ, Evans CE, Rodina A, et al. Experimental and structural testing module to analyze paralogue-specificity and affinity in the Hsp90 inhibitors series. J Med Chem. 2013;56(17):6803–18. Epub 2013/08/24. doi: 10.1021/jm400619b. PubMed PMID: 23965125; PubMed Central PMCID: PMCPMC3985615.

35. Cermak N, Olcum S, Delgado FF, Wasserman SC, Payer KR, M AM, et al. High-throughput measurement of single-cell growth rates using serial microfluidic mass sensor arrays. Nat Biotechnol. 2016;34(10):1052–9. Epub 2016/09/07. doi: 10.1038/nbt.3666. PubMed PMID: 27598230; PubMed Central PMCID: PMCPMC5064867.

36. Ngwa VM, Edwards DN, Philip M, Chen J. Microenvironmental Metabolism Regulates Antitumor Immunity. Cancer Res. 2019;79(16):4003–8. Epub 2019/08/01. doi: 10.1158/0008-5472.CAN-19-0617. PubMed PMID: 31362930; PubMed Central PMCID: PMCPMC6697577.

37. Welihinda AA, Kaur M, Raveendran KS, Amento EP. Enhancement of inosine-mediated A2AR signaling through positive allosteric modulation. Cell Signal. 2018;42:227–35. Epub 2017/11/12. doi: 10.1016/j.cellsig.2017.11.002. PubMed PMID: 29126977; PubMed Central PMCID: PMCPMC5732040.

38. Stine ZE, Walton ZE, Altman BJ, Hsieh AL, Dang CV. MYC, Metabolism, and Cancer. Cancer Discov. 2015;5(10):1024–39. Epub 2015/09/19. doi: 10.1158/2159-8290.CD-15-0507. PubMed PMID: 26382145; PubMed Central PMCID: PMCPMC4592441.

39. Nguyen L, Papenhausen P, Shao H. The Role of c-MYC in B-Cell Lymphomas: Diagnostic and Molecular Aspects. Genes (Basel). 2017;8(4). Epub 2017/04/06. doi: 10.3390/genes8040116. PubMed PMID: 28379189; PubMed Central PMCID: PMCPMC5406863.

40. Ennishi D, Jiang A, Boyle M, Collinge B, Grande BM, Ben-Neriah S, et al. Double-Hit Gene Expression Signature Defines a Distinct Subgroup of Germinal Center B-Cell-Like Diffuse Large B-Cell Lymphoma. J Clin Oncol. 2019;37(3):190–201. Epub 2018/12/14. doi: 10.1200/JCO.18.01583. PubMed PMID: 30523716; PubMed Central PMCID: PMCPMC6804880.

41. Basso K, Margolin AA, Stolovitzky G, Klein U, Dalla-Favera R, Califano A. Reverse engineering of regulatory networks in human B cells. Nat Genet. 2005;37(4):382–90. Epub 2005/03/22. doi: 10.1038/ng1532. PubMed PMID: 15778709.

42. Ellis RJ. Macromolecular crowding: obvious but underappreciated. Trends Biochem Sci. 2001;26(10):597–604. Epub 2001/10/09. PubMed PMID: 11590012.

43. Bereiter-Hahn J, Stubig C, Heymann V. Cell cycle-related changes in F-actin distribution are correlated with glycolytic activity. Experimental cell research. 1995;218(2):551–60. Epub 1995/06/01. doi: 10.1006/excr.1995.1190. PubMed PMID: 7796889.

44. Jain S, Wheeler JR, Walters RW, Agrawal A, Barsic A, Parker R. ATPase-Modulated Stress Granules Contain a Diverse Proteome and Substructure. Cell. 2016;164(3):487–98. Epub 2016/01/19. doi: 10.1016/j.cell.2015.12.038. PubMed PMID: 26777405; PubMed Central PMCID: PMCPMC4733397.

45. Wallace EW, Kear-Scott JL, Pilipenko EV, Schwartz MH, Laskowski PR, Rojek AE, et al. Reversible, Specific, Active Aggregates of Endogenous Proteins Assemble upon Heat Stress. Cell. 2015;162(6):1286–98. Epub 2015/09/12. doi: 10.1016/j.cell.2015.08.041. PubMed PMID: 26359986; PubMed Central PMCID: PMCPMC4567705.

46. Hasko G, Sitkovsky MV, Szabo C. Immunomodulatory and neuroprotective effects of inosine. Trends Pharmacol Sci. 2004;25(3):152–7. Epub 2004/03/17. doi: 10.1016/j.tips.2004.01.006. PubMed PMID: 15019271.

47. Taldone T, Zatorska D, Patel PD, Zong H, Rodina A, Ahn JH, et al. Design, synthesis, and evaluation of small molecule Hsp90 probes. Bioorg Med Chem. 2011;19(8):2603–14. Epub 2011/04/05. doi: 10.1016/j.bmc.2011.03.013. PubMed PMID: 21459002; PubMed Central PMCID: PMCPMC3143825.

48. Yamada T, Letunic I, Okuda S, Kanehisa M, Bork P. iPath2.0: interactive pathway explorer. Nucleic Acids Res. 2011;39(Web Server issue):W412–5. Epub 2011/05/07. doi: 10.1093/nar/gkr313. PubMed PMID: 21546551; PubMed Central PMCID: PMCPMC3125749.

49. Burg TP, Godin M, Knudsen SM, Shen W, Carlson G, Foster JS, et al. Weighing of biomolecules, single cells and single nanoparticles in fluid. Nature. 2007;446(7139):1066–9. Epub 2007/04/27. doi: 10.1038/nature05741. PubMed PMID: 17460669.

50. Zhang B, Kirov S, Snoddy J. WebGestalt: an integrated system for exploring gene sets in various biological contexts. Nucleic Acids Res. 2005;33(Web Server issue):W741–8. Epub 2005/06/28. doi: 10.1093/nar/gki475. PubMed PMID: 15980575; PubMed Central PMCID: PMCPMC1160236.

