## supplementary information for "HSP90 facilitates oncogenic alterations of metabolism in B-cell lymphomas"

#### SUPPLEMENTAL TABLES

**TABLE S1.** KEGG pathway enrichment analysis of the cytoplasmic HSP90 interactome

**TABLE S2.** STRING-based categorization of the subset of metabolic HSP90 interactome.

**TABLE S3.** Exo-metabolomics analysis of 50 DLBCL patients vs. 25 age- and sex-matched healthy individuals.

**TABLE S4.** Spearman's' rank correlation of MYC expression with the subset of transcripts corresponding to the HSP90 metabolic interactome proteins in two patient cohorts (WCM and TCGA) and one cell line cohort.

#### SUPPLEMENTAL FIGURES

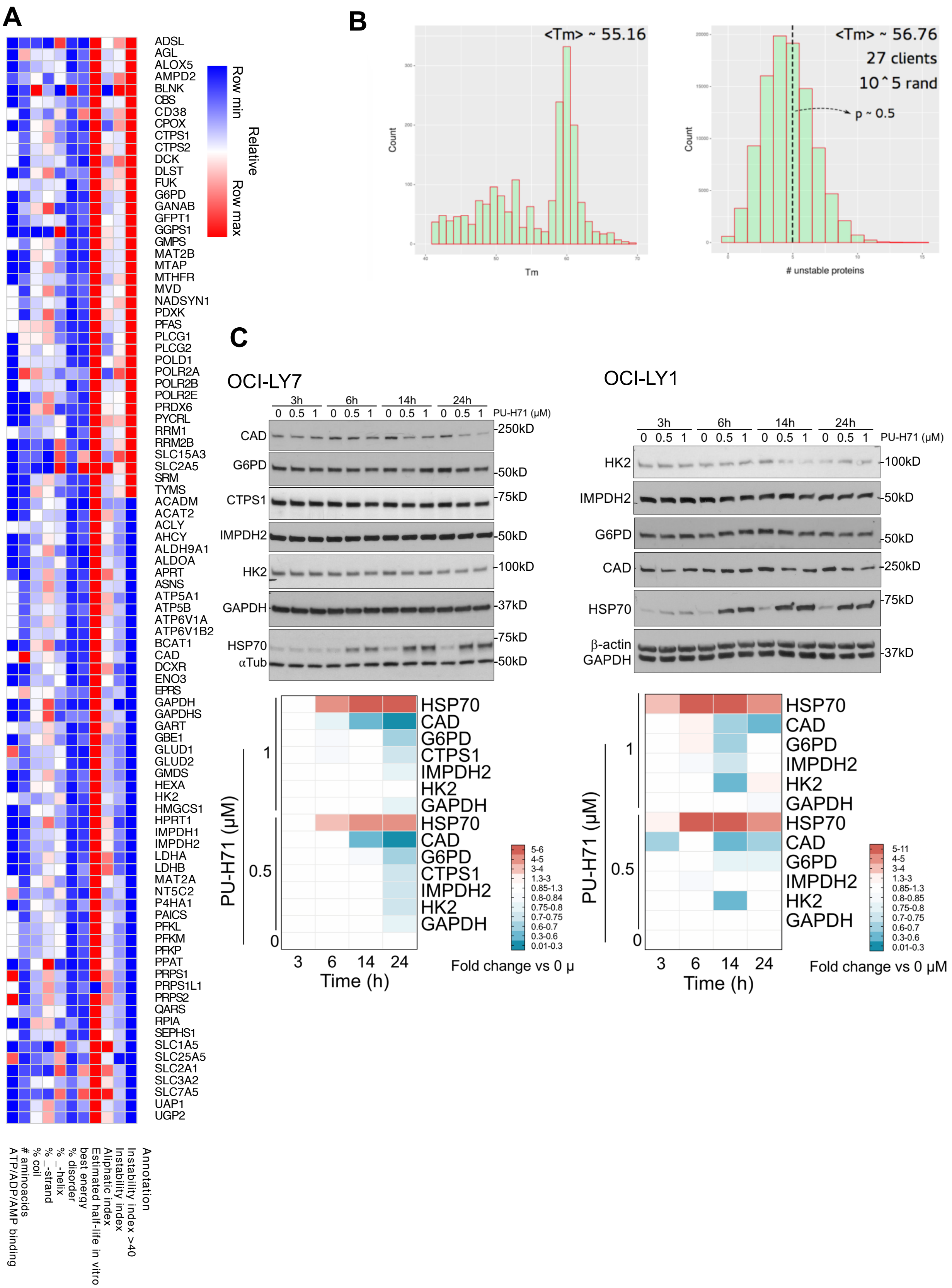
A
ADSL
AGL
ALOX5
AMPD2
BLNK
CBS
CD38
CPOX
CTPS1
CTPS2
DCK
DLST
FUK
G6PD
GANAB
GFPT1
GGPS1
GMPS
MAT2B
MTAP
MTHFR
MVD
NADSYN1
PDXK
PFAS
PLCG1
PLCG2
POLD1
POLR2A
POLR2B
POLR2E
PRDX6
PYCRL
RRM1
RRM2B
SLC15A3
SLC2A5
SRM
TYMS
ACADM
ACAT2
ACLY
AHCY
ALDH9A1
ALDOA
APRT
ASNS
ATP5A1
ATP5B
ATP6V1A
ATP6V1B2
BCAT1
CAD
DCXR
ENO3
EPRS
GAPDH
GAPDHS
GART
GBE1
GLUD1
GLUD2
GMDS
HEXA
HK2
HMGCS1
HPRT1
IMPDH1
IMPDH2
LDHA
LDHB
MAT2A
NT5C2
P4HA1
PAICS
PFKL
PFKM
PFKP
PPAT
PRPS1
PRPS1L1
PRPS2
QARS
RPIA
SEPHS1
SLC1A5
SLC25A5
SLC2A1
SLC3A2
SLC7A5
UAP1
UGP2
Row min
Relative
Row max
ATP/ADP/AMP binding
### aminoacids
% coil
% _strand
% _helix
% disorder
best energy
Estimated half-life in vitro
Aliphatic index
Instability index
Instability index >40
Annotation
B
<Tm> ~ 55.16
Count
Tm
<Tm> ~ 56.76
27 clients
10^5 rand
p ~ 0.5
Count
### unstable proteins
C
OCI-LY7
3h 6h 14h 24h
0 0.5 1 0 0.5 1 0 0.5 1 0 0.5 1 PU-H71 (µM)
CAD -250kD
G6PD -50kD
CTPS1 -75kD
IMPDH2 -50kD
HK2 -100kD
GAPDH -37kD
HSP70 -75kD
αTub -50kD
OCI-LY1
3h 6h 14h 24h
0 0.5 1 0 0.5 1 0 0.5 1 0 0.5 1 PU-H71 (µM)
HK2 -100kD
IMPDH2 -50kD
G6PD -50kD
CAD -250kD
HSP70 -75kD
β-actin
GAPDH -37kD
PU-H71 (µM)
1
0.5
0
HSP70
CAD
G6PD
CTPS1
IMPDH2
HK2
GAPDH
HSP70
CAD
G6PD
CTPS1
IMPDH2
HK2
GAPDH
3 6 14 24
Time (h)
5-6
4-5
3-4
1.3-3
0.85-1.3
0.8-0.84
0.75-0.8
0.7-0.75
0.6-0.7
0.3-0.6
0.01-0.3
Fold change vs 0 µ
PU-H71 (µM)
1
0.5
0
HSP70
CAD
G6PD
IMPDH2
HK2
GAPDH
HSP70
CAD
G6PD
IMPDH2
HK2
GAPDH
3 6 14 24
Time (h)
5-11
4-5
3-4
1.3-3
0.85-1.3
0.8-0.85
0.75-0.8
0.7-0.75
0.6-0.7
0.3-0.6
0.01-0.3
Fold change vs 0 µM

**Figure S1. Biochemical properties of the proteins included in the HSP90 metabolic interactome. A:** HSP90 metabolic interactome components are depicted in rows. Biochemical properties of proteins are shown on the bottom. Heat map represents maximum and minimum properties for this set. **B:** ***left:*** Histogram of Tm values (mammalian cells) extracted from "Leuenberger P, et al. Cell-wide analysis of protein thermal unfolding reveals determinants of thermostability. Science 2017;355,6327,eaai7825". Mean Tm (<Tm>) value was found to be ~ 55.16 C. ***right:*** Tm values for a sub-set of 27 HSP90 metabolic interactome proteins (<Tm> ~ 56.76 C). Five proteins in this sub-set were found to be described as 'unstable' (i.e., Tm < 47.92 C). A randomization analysis (n = $10^5$) was performed to determine if this subset was enriched in unstable proteins. **C:** Effect of long-term HSP90 inhibition on the levels of metabolic protein cargoes**.** Time course (3, 6, 14 and 24 h) of the abundance of HSP90 metabolic interactome components CAD, G6PD, CTPS1, IMPDH2, HK2 and GAPSH upon HSP90 inhibition with increasing concentrations of PU-H71 (0.5 and 1 µM) or vehicle (0) in OCI-LY7 and OCI-LY1 cells. HSP70 was used as molecular readout of effective HSP90 inhibition. Densitometry of blots are shown at the bottom as color-coded fold changes over vehicle control.

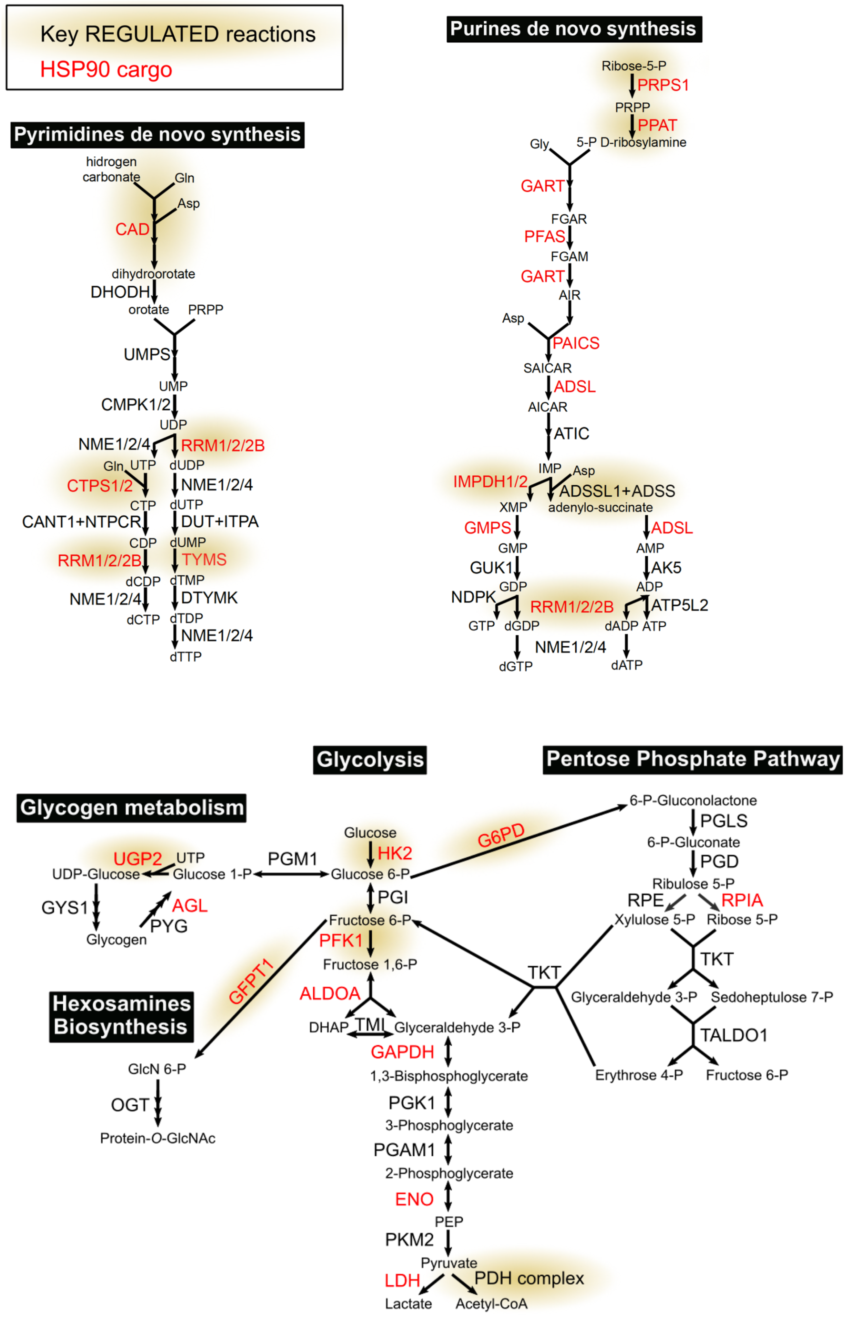


**Figure S2.** Nucleotides and carbohydrates pathways indicating key regulated reactions with yellow shadowing and enzymes from the metabolic HSP90 intreactome in red.

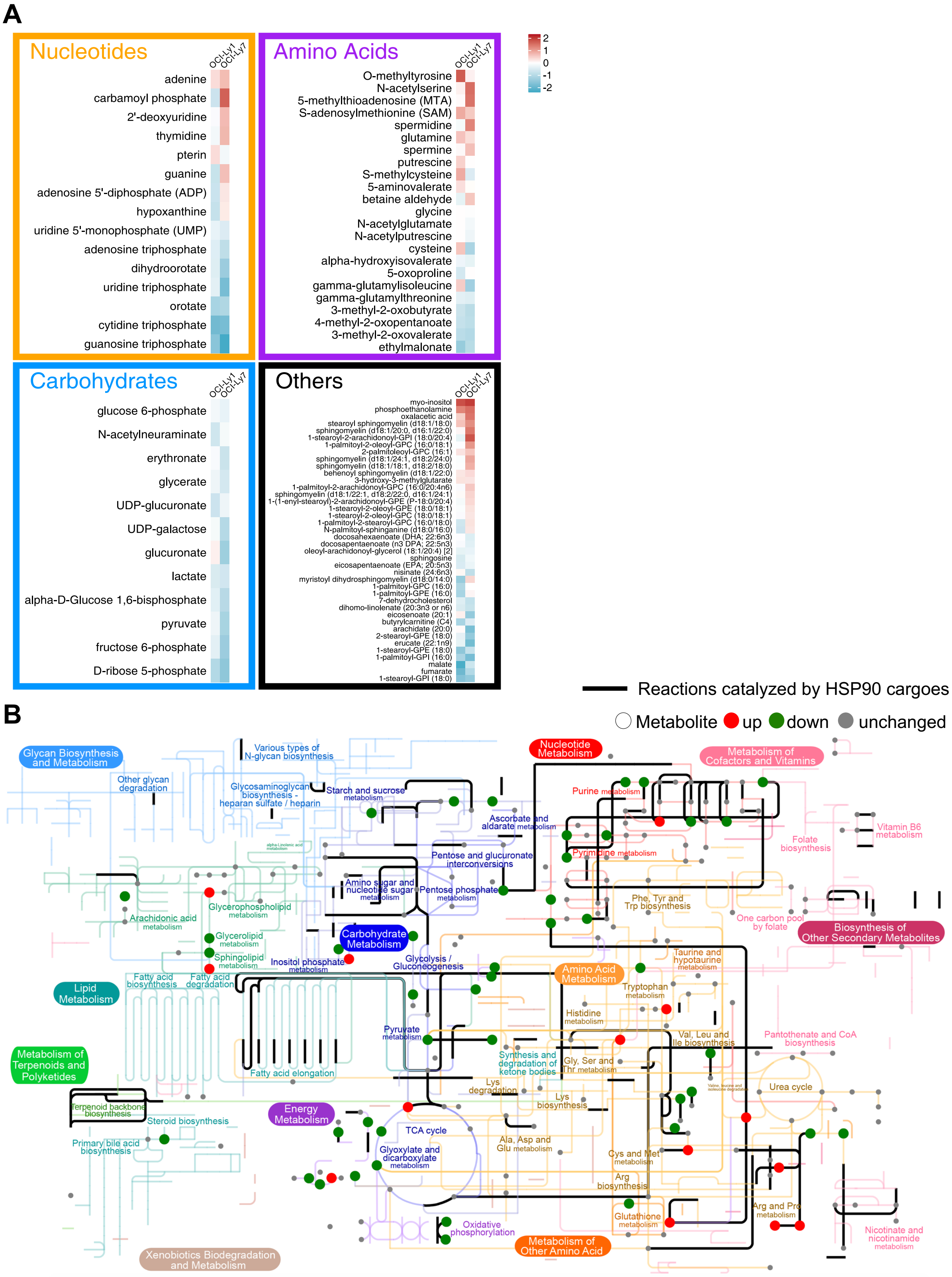
A
Nucleotides
OCI-Ly1
OCI-Ly7
adenine
carbamoyl phosphate
2'-deoxyuridine
thymidine
pterin
guanine
adenosine 5'-diphosphate (ADP)
hypoxanthine
uridine 5'-monophosphate (UMP)
adenosine triphosphate
dihydroorotate
uridine triphosphate
orotate
cytidine triphosphate
guanosine triphosphate
Amino Acids
O-methyltyrosine
N-acetylserine
5-methylthioadenosine (MTA)
S-adenosylmethionine (SAM)
spermidine
glutamine
spermine
putrescine
S-methylcysteine
5-aminovalerate
betaine aldehyde
glycine
N-acetylglutamate
N-acetylputrescine
cysteine
alpha-hydroxyisovalerate
5-oxoproline
gamma-glutamylisoleucine
gamma-glutamylthreonine
3-methyl-2-oxobutyrate
4-methyl-2-oxopentanoate
3-methyl-2-oxovalerate
ethylmalonate
2
1
0
-1
-2
Carbohydrates
glucose 6-phosphate
N-acetylneuraminate
erythronate
glycerate
UDP-glucuronate
UDP-galactose
glucuronate
lactate
alpha-D-Glucose 1,6-bisphosphate
pyruvate
fructose 6-phosphate
D-ribose 5-phosphate
Others
myo-inositol
phosphoethanolamine
oxalacetic acid
stearoyl sphingomyelin (d18:1/18:0)
sphingomyelin (d18:1/20:0, d16:1/22:0)
1-stearoyl-2-arachidonoyl-GPI (18:0/20:4)
1-palmitoyl-2-oleoyl-GPC (16:0/18:1)
2-palmitoleoyl-GPC (16:1)
sphingomyelin (d18:1/24:1, d18:2/24:0)
sphingomyelin (d18:1/18:1, d18:2/18:0)
behenoyl sphingomyelin (d18:1/22:0)
3-hydroxy-3-methylglutarate
1-palmitoyl-2-arachidonoyl-GPC (16:0/20:4n6)
sphingomyelin (d18:1/22:1, d18:2/22:0, d16:1/24:1)
1-(1-enyl-stearoyl)-2-arachidonoyl-GPE (P-18:0/20:4)
1-stearoyl-2-oleoyl-GPE (18:0/18:1)
1-stearoyl-2-oleoyl-GPC (18:0/18:1)
1-palmitoyl-2-stearoyl-GPC (16:0/18:0)
N-palmitoyl-sphinganine (d18:0/16:0)
docosahexaenoate (DHA; 22:6n3)
docosapentaenoate (n3 DPA; 22:5n3)
oleoyl-arachidonoyl-glycerol (18:1/20:4) [2]
sphingosine
eicosapentaenoate (EPA; 20:5n3)
nisinate (24:6n3)
myristoyl dihydrosphingomyelin (d18:0/14:0)
1-palmitoyl-GPC (16:0)
1-palmitoyl-GPE (16:0)
7-dehydrocholesterol
dihomo-linolenate (20:3n3 or n6)
eicosenoate (20:1)
butyrylcarnitine (C4)
arachidate (20:0)
2-stearoyl-GPE (18:0)
erucate (22:1n9)
1-stearoyl-GPE (18:0)
1-palmitoyl-GPI (16:0)
malate
fumarate
1-stearoyl-GPI (18:0)
B
Reactions catalyzed by HSP90 cargoes
Metabolite up down unchanged
Glycan Biosynthesis and Metabolism
Various types of N-glycan biosynthesis
Other glycan degradation
Glycosaminoglycan biosynthesis - heparan sulfate / heparin
Starch and sucrose metabolism
Nucleotide Metabolism
Metabolism of Cofactors and Vitamins
Purine metabolism
Ascorbate and aldarate metabolism
Vitamin B6 metabolism
Folate biosynthesis
Pyrimidine metabolism
Pentose and glucuronate interconversions
Amino sugar and nucleotide sugar metabolism
Pentose phosphate metabolism
Phe, Tyr and Trp biosynthesis
One carbon pool by folate
Biosynthesis of Other Secondary Metabolites
Arachidonic acid metabolism
Glycerophospholipid metabolism
Glycerolipid metabolism
Sphingolipid metabolism
Carbohydrate Metabolism
Inositol phosphate metabolism
Glycolysis / Gluconeogenesis
Taurine and hypotaurine metabolism
Amino Acid Metabolism
Tryptophan metabolism
Lipid Metabolism
Fatty acid biosynthesis
Fatty acid degradation
Histidine metabolism
Pyruvate metabolism
Val, Leu and Ile biosynthesis
Pantothenate and CoA biosynthesis
Metabolism of Terpenoids and Polyketides
Synthesis and degradation of ketone bodies
Gly, Ser and Thr metabolism
Fatty acid elongation
Urea cycle
Lys degradation
Lys biosynthesis
Terpenoid backbone biosynthesis
Energy Metabolism
Steroid biosynthesis
TCA cycle
Primary bile acid biosynthesis
Ala, Asp and Glu metabolism
Glyoxylate and dicarboxylate metabolism
Cys and Met metabolism
Arg biosynthesis
Glutathione metabolism
Arg and Pro metabolism
Oxidative phosphorylation
Metabolism of Other Amino Acid
Nicotinate and nicotinamide metabolism
Xenobiotics Biodegradation and Metabolism

**Figure S3. HSP90 metabolome in lymphoma cells. A:** Heatmap of differentially ($p<0.05$) changed metabolites in OCI-LY1 and OCI-LY7 cells treated with PU-H71 for 6 h. **B:** Overlay of metabolic enzymes from the HSP90 interactome and PU-H71-modified cellular metabolites onto KEGG human metabolism map. Metabolites are represented by red (up with treatment), green (down) and grey (unchanged) circles. Reactions catalyzed by HSP90 metabolic cargoes are shown as black lines. Metabolic pathways are represented in KEGG default colors.

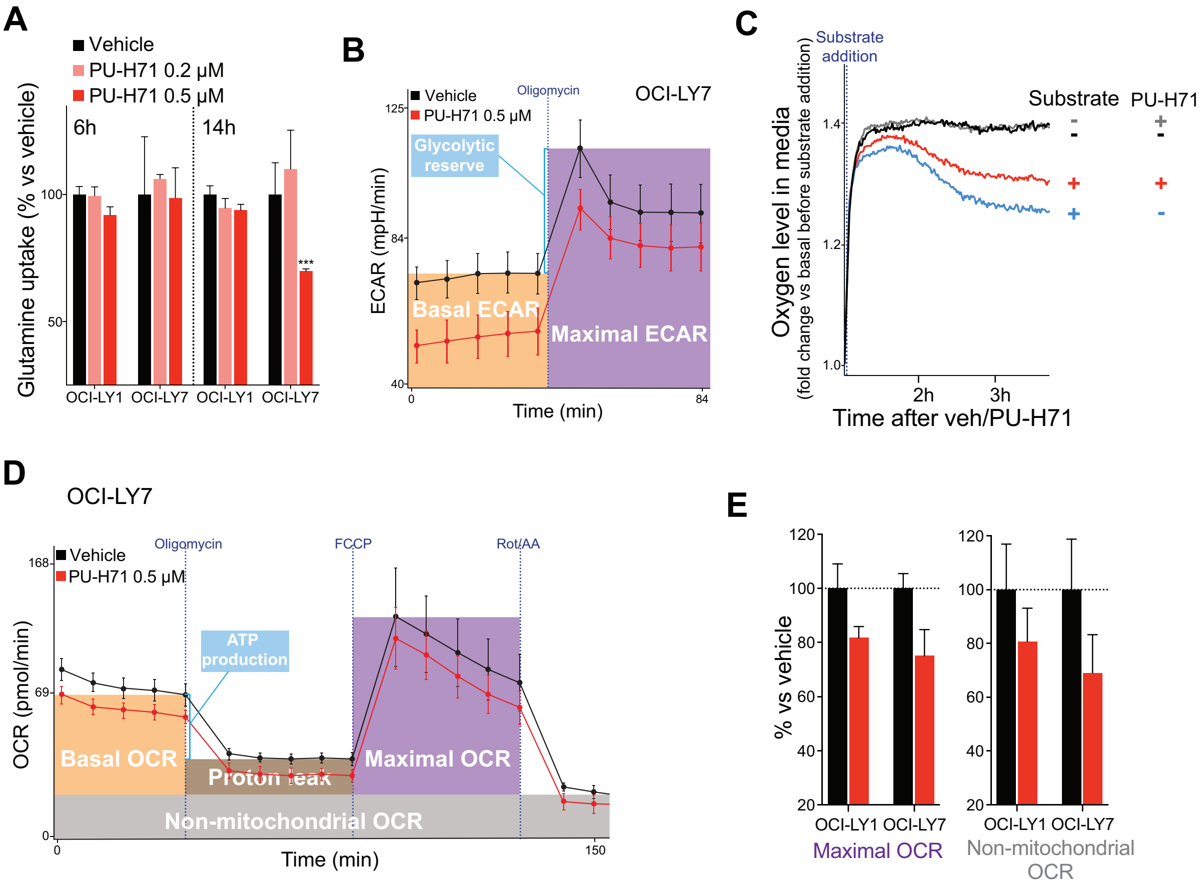


**Figure S4. Effect of HSP90 inhibition on glutamine uptake, glycolysis and oxygen consumption. A:** Glutamine uptake in OCI-LY1 and OCI-LY7 cells treated with vehicle (black), PU-H71 0.2 µM (pink) and PU-H71 0.5 µM (red) for 6 h and 14 h. Data is normalized to vehicle-treated cells. **B:** Extracellular acidification rate (ECAR) of OCI-LY7 cells treated with vehicle (black) or PU-H71 (red), at baseline and upon oligomycin treatment to estimate the maximal ECAR. Error bars are SD of 10 replicate wells. Representative experiment of triplicates shown. **C:** Mitochondrial respiration as determined by real-time measurement of oxygen levels in the tissue culture medium of BL Raji cells in absence of glutamine with vehicle (black) or PU-H71 (grey), and presence of glutamine with vehicle (blue) or PU-H71 (red). Representative experiment of triplicates shown. **D:** Oxygen consumption rate (OCR) of OCI-LY7 cells treated with vehicle (black) or PU-H71 (red) at baseline, upon oligomycin treatment to determine proton leak and OCR-linked ATP production, upon FCCP to estimate maximal OCR, and upon rotenone/antimycin A to estimate non-mitochondrial OCR. **E:** Maximal OCR and non-mitochondrial OCR in OCI-LY1 and OCI-LY7 cells treated with vehicle (black) or PU-H71 (red). In all panels, unless stated differently, error bars are SEM of 3 independent experiments. p values were calculated by T-test. n.s., not significant, *<0.05, **<0.01 and ***<0.001.

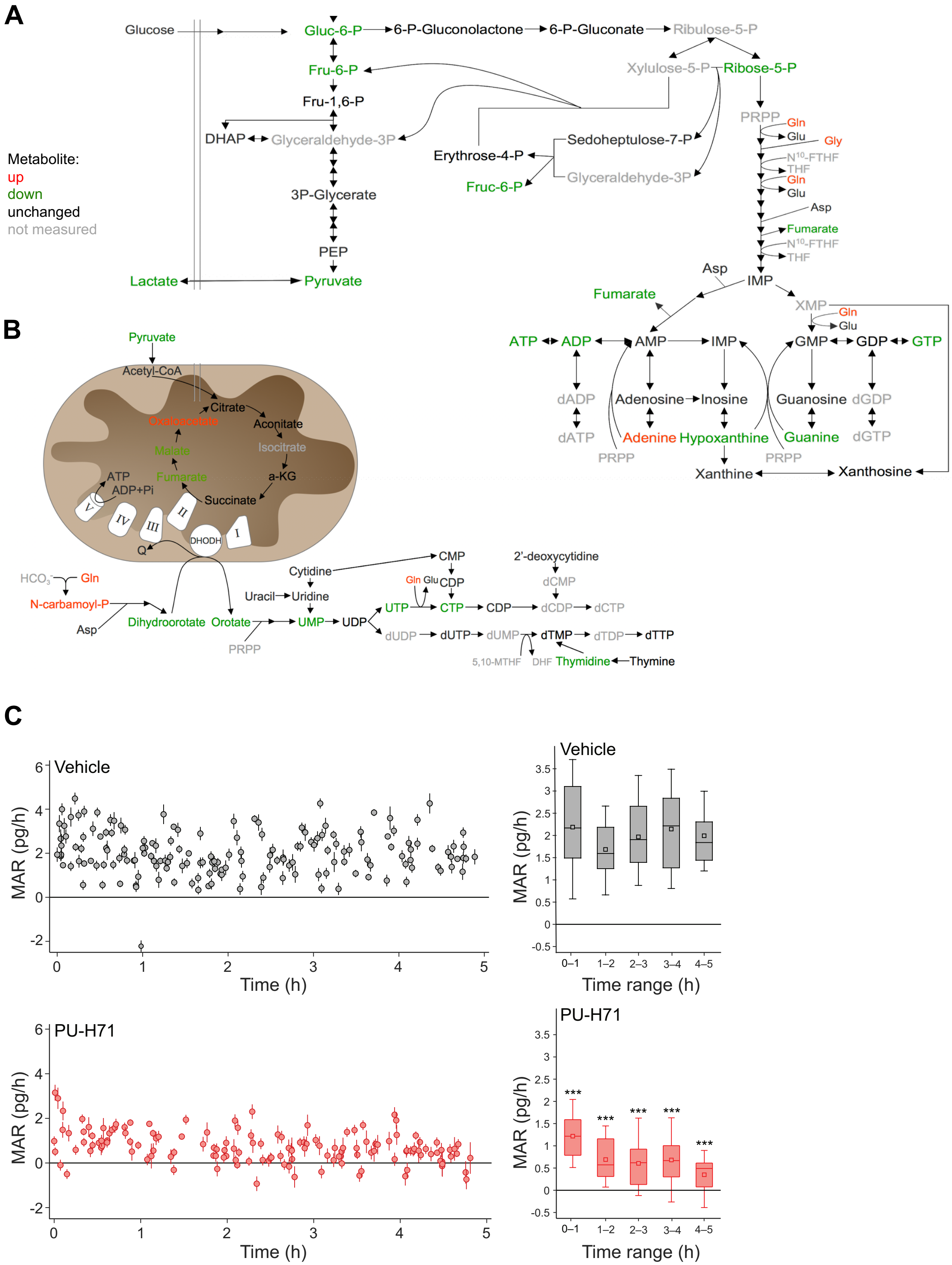
A
Glucose
Gluc-6-P
6-P-Gluconolactone
6-P-Gluconate
Ribulose-5-P
Xylulose-5-P
Ribose-5-P
Fru-6-P
Fru-1,6-P
DHAP
Glyceraldehyde-3P
Sedoheptulose-7-P
Erythrose-4-P
Fruc-6-P
3P-Glycerate
PEP
Lactate
Pyruvate
Metabolite:
up
down
unchanged
not measured
PRPP
Gln
Glu
Gly
N10-FTHF
THF
Asp
Fumarate
IMP
XMP
ATP
ADP
AMP
GMP
GDP
GTP
dADP
dATP
Adenosine
Inosine
Guanosine
dGDP
dGTP
Adenine
Hypoxanthine
Guanine
Xanthine
Xanthosine
B
Pyruvate
Acetyl-CoA
Citrate
Oxaloacetate
Aconitate
Isocitrate
Malate
Fumarate
a-KG
Succinate
ATP
ADP+Pi
V
IV
III
II
I
DHODH
Q
HCO3
N-carbamoyl-P
Asp
Dihydroorotate
Orotate
PRPP
Cytidine
Uracil
Uridine
UMP
UDP
UTP
CTP
CMP
CDP
dCDP
dCTP
2'-deoxycytidine
dCMP
dUDP
dUTP
dUMP
dTMP
dTDP
dTTP
5,10-MTHF
DHF
Thymidine
Thymine
C
Vehicle
MAR (pg/h)
Time (h)
Time range (h)
0–1
1–2
2–3
3–4
4–5
PU-H71
***

**Figure S5. Metabolic and biomass changes caused by HSP90 inhibition in DLBCL cells.** Common changes in metabolites levels upon treatment of OCI-LY1 and OCI-LY7 cell lines with PU-H71 (vs. vehicle) for 6 h. Pathways represent glycolysis and PPP (**A**) and mitochondria-orotate metabolites (**B**). **C:** Real-time assessment at single-cell resolution of cellular Mass Accumulation Rate (MAR) in pg/h in OCI-LY1 cells treated with vehicle (black) or PU-H71 1 µM (red). On the right, mean MAR comparing binned datasets (from time 0 to 1, from 1 to 2, and so on).

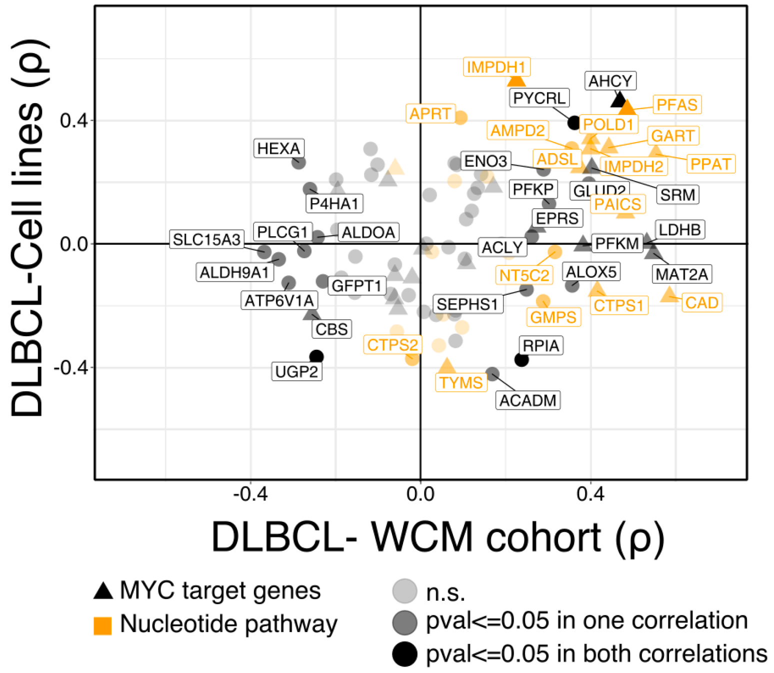


**Figure S6. Correlation of MYC and HSP90 metabolic interactome genes in DLBCL cell lines.** Spearman's rank correlation plots of gene expression of MYC and HSP90 metabolic interactome components in the WCM cohort of DLBCL patients (n = 80) vs. a cohort of 30 DLBCL cell lines. Canonical MYC target genes are depicted with triangles. Genes from the nucleotide pathway are shown in yellow. Darker colors indicate significant (p<0.05) correlations in both cohorts.
